# Minimizing Reference Bias with an Impute-First Approach

**DOI:** 10.1101/2023.11.30.568362

**Authors:** Kavya Vaddadi, Taher Mun, Ben Langmead

## Abstract

Pangenome indexes reduce reference bias in sequencing data analysis. However, bias can be reduced further by using a personalized reference, e.g. a diploid human reference constructed to match a donor individual’s alleles. We present a novel impute-first alignment framework that combines elements of genotype imputation and pangenome alignment. It begins by genotyping the individual using only a subsample of the input reads. It next uses a reference panel and efficient imputation algorithm to impute a personalized diploid reference. Finally, it indexes the personalized reference and applies a read aligner, which could be a linear or graph aligner, to align the full read set to the personalized reference. This framework achieves higher variant-calling recall (99.54% vs. 99.37%), precision (99.36% vs. 99.18%), and F1 (99.45% vs. 99.28%) compared to a graph pangenome aligner. The personalized reference is also smaller and faster to query compared to a pangenome index, making it an overall advantageous choice for whole-genome DNA sequencing experiments.

## 1 Introduction

The field of Bioinformatics has developed various ways to reduce the “reference bias” that results when aligning reads to reference genomes [1]. Reference bias occurs when reads containing non-reference alleles fail to align to their true point of origin, leading to inaccurate results for analyses concerned with hypervariable regions [2], allele-specific effects [3–6], ancient DNA [7, 8] or epigenomics [9]. Pangenome alignment methods reduce bias by using a pangenome graph [10–12] or collection of linear genomes [13]. Pangenome indexes allow the aligner to remove unwanted penalties associated with known genetic variants. While these methods reduce bias [14–16], personalized references, which are tailored to include the specific alleles present in the individual under study (the “donor”) are known to be better at reducing bias [15, 16].

Here we present an efficient “impute-first alignment” framework combining advantages of genotype imputation and pangenome alignment. Modern imputation tools are efficient [17, 18] and leverage comprehensive genetic and linkage-disequilibrium information from reference panels like the 1000 Genomes Project (1KGP) [19], which spans 1000s of haplotypes, and Human Genome Structural Variation Consortium (HGSVC2) [20], which includes complex structural variations. A key aspect of these panels is that they are phased, meaning they faithfully represent the co-inheritance patterns of nearby variants.

Our framework begins by taking a subsample of the input reads and performing genotyping and imputation steps. The output of this “personalization” process is a diploid personalized reference. We then index the personalized reference and align all the input reads to the personalized index. Here we use a pangenome aligner, where the “pangenome” contains exactly the alleles imputed for the donor. This reduces reference bias while also reducing computational overhead relative to methods that use a large non-personalized pangenome.

Past methods used personalized references to alleviate reference bias in the study of RNA-seq [5, 21, 22], but rely on the user to provide the donor’s genotypes as an input. Some methods additionally perform a personalization step, but have other drawbacks. Gramtools [23] builds a pangenome reference representation that can be used both to align reads and to perform imputation of a personalized genome. That study was limited to monoploids and showed that a personalized genome for Plasmodium falciparum helps to alleviate reference bias in hypervariable regions. iCORN [24] gradually refines the reference to contain more ALT alleles through an expensive iterative alignment process, and also focuses on monoploids. MMSeq [25] performs variant calling on an input BAM file to construct a personalized diploid transcriptome, but does not perform imputation. RefEditor [26] uses a genotype imputation tool (MaCH or Minimac) to obtain phased diploid genotypes based on SNP genotyping array data. By contrast, our work performs personalization based on the input (sequencing) dataset itself and does not require a second dataset. Importantly, none of the above tools used an efficient imputation method like Glimpse [18], which has the positional Burrows Wheeler Transform (PBWT) [27] as its computational engine. Further, none of the above-mentioned studies assessed the computational cost, and thus the practicality, of an efficient personalization phase that runs as part of the main data analysis.

Groza et al. [9] introduced Graph Personal Genome, or a graph-shaped reference containing only the donor’s diploid variants. In their case, use of a personalized reference is a first step toward obtaining less biased ChIP-seq peak calls. Our framework also uses this idea. However, our framework both imputes the diploid variants and creates this graph-genome reference as part of the same data analysis. The Groza et al. framework takes the variants as an input, with the expectation that these would be called from a separate dataset. Also unlike Groza et al. we compare the method of constructing a personalized graph-genome reference to the more common practice of simply using a pangenome reference built over a large panel, i.e. the 1000 Genomes Project phase-3 callset.

We show that our impute-first framework can construct a diploid personalized reference at a low computational cost relative to the indexing of a pangenome. Further, we show that these personalized references are accurate, having high variant-level precision and recall, and accurate overall phasing. Finally, we show that a pangenome index built over a personalized genome is more computationally efficient, less biased, and produces more accurate downstream variant calls compared to a pangenome index, making the impute-first framework an appealing choice for whole-genome DNA sequencing data analyses.

## 2 Results

### 2.1 Overview of Impute-first Alignment Workflow

Figure 1 illustrates the workflow, which is divided into two components. In the first, a sample of the input reads is analyzed using a pangenome genotyper (e.g. Rowbowt [34]) and imputation tool (e.g. Glimpse [18]). The output is a phased personalized variant callset, which in turn is used to construct a personalized diploid reference genome. In the second component, the personalized reference is indexed (e.g. using VG Giraffe) and a read aligner analyzes the full read set with respect to this index. The workflow is modular; different tools can be substituted for the initial genotyping step (e.g. Bowtie2+bcftools instead of Rowbowt), the imputation step (e.g. Beagle instead of Glimpse) and the final read alignment step (e.g. Bowtie2 or BWA-MEM instead of VG Giraffe). The table uses underlining to highlight short “designators” that we use later to refer to particular tools, e.g. “r” for Rowbowt and “bb” for Bowtie2+bcftools.

**Figure 1:**
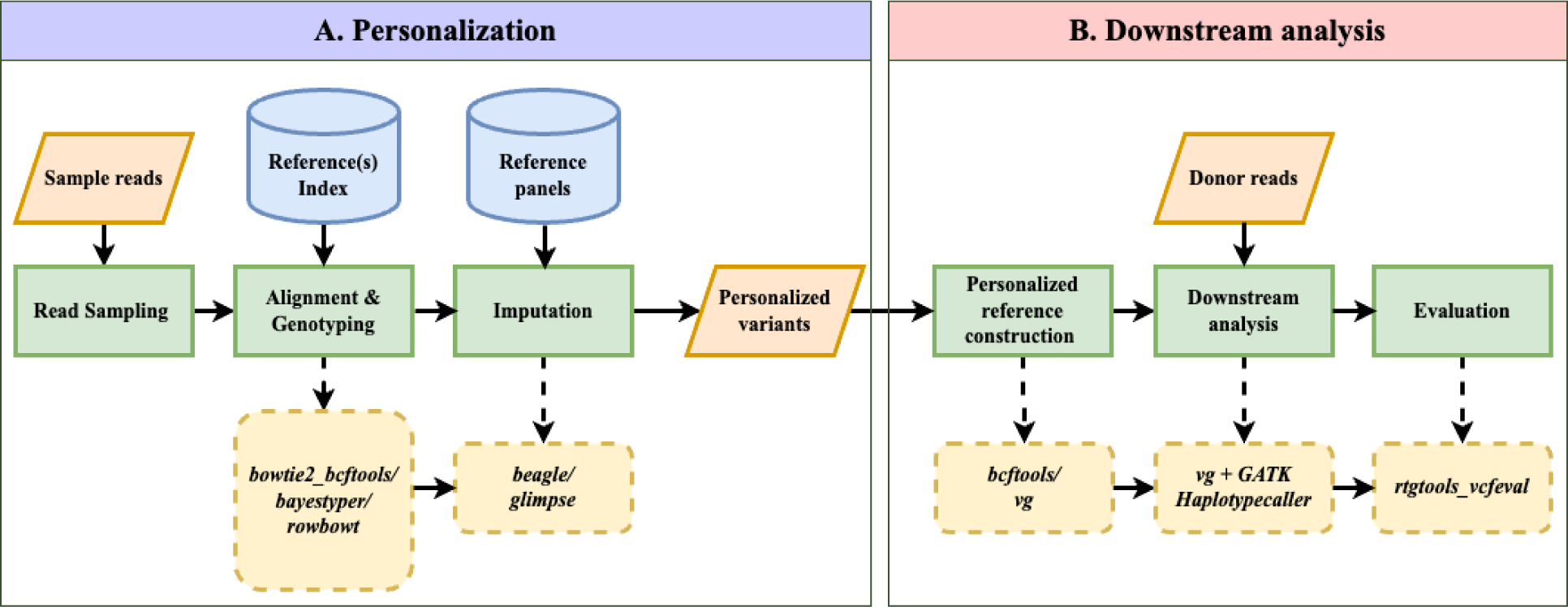
Impute-first alignment workflow in the context of analyzing a human whole-genome sequencing dataset. Box A shows the workflow up until it creates a personalized diploid reference. This the “personalization” component. Box B continues the workflow by aligning the full set of reads to the personalized reference. This is the “downstream” component.

### 2.2 Personalization

We studied personalization workflows that used various algorithms and reference representations. The Bowtie2+bcftools workflow used a linear reference index, Rowbowt used a collection of linear haplotypes indexed with *r*-index, and BayesTyper used a graph pangenome index. We evaluated the workflows’ abilities to efficiently call correct phased genotypes. We tested the workflows on a subsample of reads taken from each of two whole-genome DNA sequencing datasets. The first dataset was derived from individual HG001/NA12878 and was sequenced as part of the 1000 Genomes Project [19]. The second was derived from individual HG002/NA24385 and was sequenced as part of the Google Brain project [28]. The initial sequencing depth for both was about 30*×*, which we subsampled to lower coverage levels ranging from 0.01*×* to 5*×* respectively (Table 1).

**Table 1:** The steps, inputs, outputs and tools used in the tested Impute-first workflows.

| Personalization |  |  |  |
| --- | --- | --- | --- |
| Step | Input | Tool (designator) | Output |
| Read Sampling | **Donor reads:** Whole-genome DNA-seq reads from HG001/NA12878 [19] & HG002/NA24385 [28] | seqtk [29] | **Reads** sampled to 0.01, 0.05, 0.1, 0.2, 0.5, 1, 2, and 5× average coverage |
| Alignment & Genotyping | **Genotype panel:** HGSVC2 [20] VCF, excluding respective HG001/NA12878 & HG002/NA24385 and family members.<br>**Reference:** GRCh38 primary assembly [30]<br>**Reads:** Output from Sampling step | bowtie2[31] + bcftools (<u>bb</u>) [32];<br>bayestyper[33] (<u>bt</u>);<br>rowbowt[34] (<u>r</u>); | **Rough genotype calls** in VCF format |
| Imputation | **Imputation panel & reference:** Same as previous step<br>**Genotype calls:** Output from genotyping step. | beagle[17] (<u>b</u>);<br>glimpse[18] (<u>g</u>) | **Personalized reference** as phased VCF file |
| Downstream analysis |  |  |  |
| Step | Input | Tool | Output |
| Personalized Reference Construction | **Personalized reference:** from Imputation step<br>**Reference:** GRCh38 primary assembly [30] | bcftools;<br>vg autoindex [11] | **Personalized reference** as diploid FASTA or graph |
| Downstream Analysis | **Personalized reference:** From above step<br>**Donor reads:** HG001/NA12878 & HG002/NA24385 [28] | vg giraffe [11]<br>gatk HaplotypeCaller [35] | **Variant calls** as VCF |
| Evaluation | **True variants:** HG001 & HG002 VCF from GIAB [36];<br>High-confidence region annotations, etc. | rtgtools<br>vcfeval [37] | **Benchmarking metrics** |

Using subsampled read sets allowed us to measure the workflow’s robustness to smaller inputs, which is relevant when (a) the input dataset is small to begin with, or (b) we are seeking to minimize time and memory taken by the personalization steps by running on only a fraction of the full input. We hypothesized that we could still achieve high personalization accuracy on subsamples, consistent with a previous study [38]. Ground-truth genotypes for both HG001 and HG002 were taken from the HGSVC2 reference panel [20, 39]. We measured different personalization workflows’ ability to correctly call alternate alleles (ALTs) and heterozygous variants (HETs). We also measured window accuracy, i.e. the fraction of 200 base-pair windows in the imputed diploid genome that exactly match the same window in the true genome.

#### Call Accuracy

We assessed genotype accuracy across various coverages both before and after imputation. We measured precision, recall, and F1 score, i.e. the harmonic mean of precision and recall.

For both HG001 and HG002, increased coverage led to more accurate calls (Figures 2 and S5). Also, imputation led to notable improvements in F1 for both ALT and HET calls for all three genotyping methods. For all three genotyping methods, Glimpse produced higher F1 for HET calls than Beagle.

**Figure 2:**
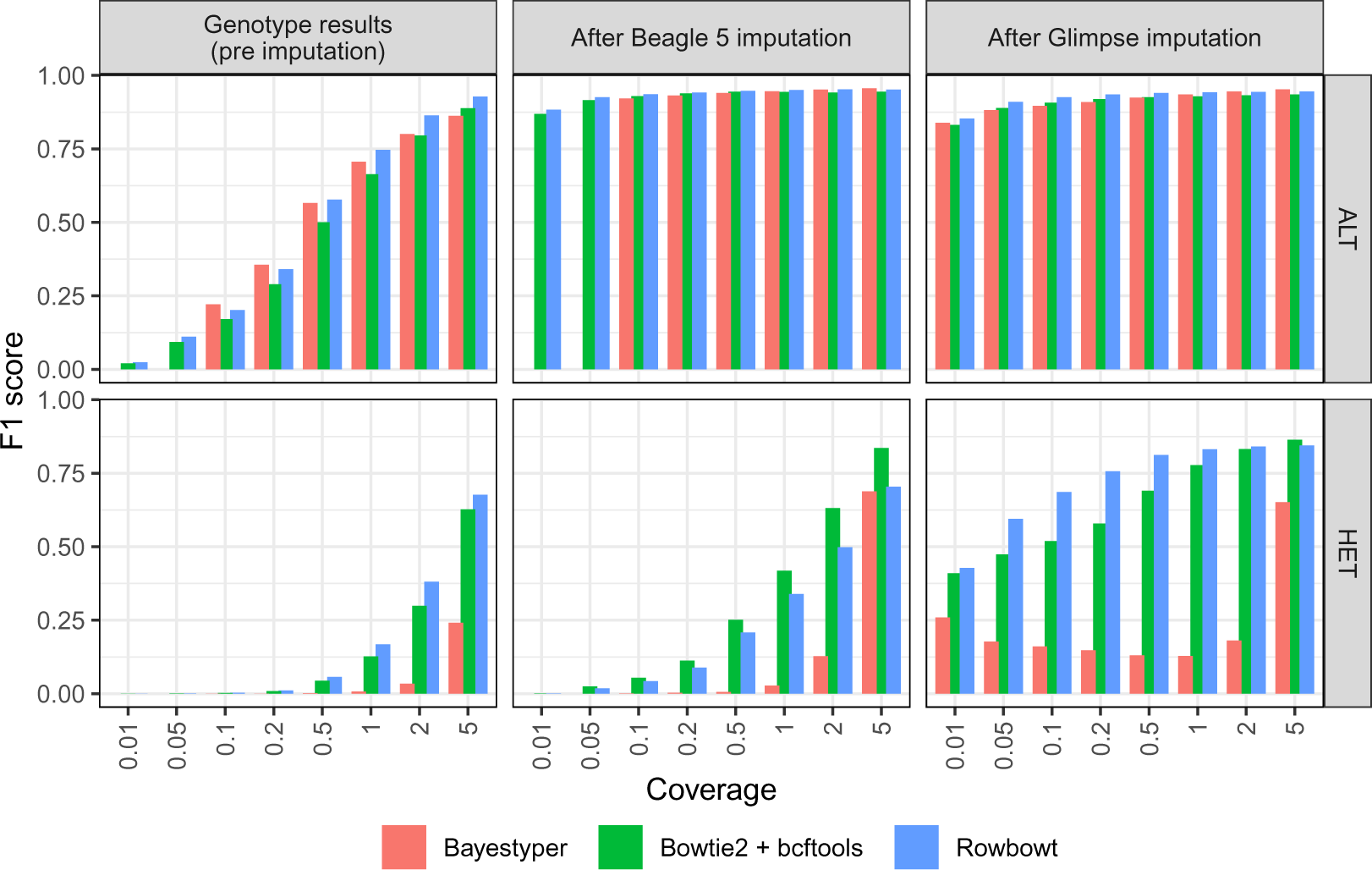
Aggregate F1 scores of alternate allele calls (ALT) and heterozygous calls (HET) across all variant types, generated using each alignment/genotyping method in the Impute-first alignment workflow.

In most scenarios, Rowbowt produced more accurate genotypes, both before and after imputation, compared to the other methods. An exception was the HET variants as imputed by Beagle, where the Bowtie2+bcftools workflow achieved somewhat higher F1. BayesTyper generally produced less accurate genotypes and imputed genotypes compared to the other tools, though this gap closed somewhat at higher coverages. For HG002, BayesTyper performed somewhat better for ALT calls, but still underperformed for HET calls (Figure S5).

#### Window Accuracy

Window accuracy is a measure of how frequently a window of genomic bases covering one or more polymorphic sites is correctly inferred in the personalized reference. We considered each of the 18.8M sites that were polymorphic according to the HGSVC2 reference panel. We call the site under consideration the “pivot.” As a group, we consider the pivot as well as all other polymorphic sites to the left of and within 200 bp of the pivot. For each such group, we determined whether all variant calls for all variants in the group were correctly genotyped and correctly phased. Figure 3 illustrates the results for HG001, with windows stratified according to the total number of polymorphic sites falling inside. The “1 – 5” stratum includes groups with 1–5 polymorphic sites, “6 – 10” stratum includes groups with 6–10 polymorphic sites, and “11+” includes groups with 11 or more polymorphic sites. Similarly, Figure S6 presents the results for HG002 using the same stratification criteria. In both cases, as expected, window accuracy increases with average coverage, since more total evidence leads to more accurate genotyping calls. Also as expected, window accuracy decreases as we consider denser strata (11+ being the densest), reflecting the increased difficulty of correctly calling all phased genotypes for all variants in a large group.

Across both HG001 and HG002, Rowbowt generally achieved the highest overall fraction of correctly-called windows. Rowbowt performed particularly well at *≤* 1x coverage. Conversely, in regions with *>* 1x coverage, when used in combination with Beagle, Bowtie2+bcftools achieved higher (sometimes highest overall) accuracy.

**Figure 3:**
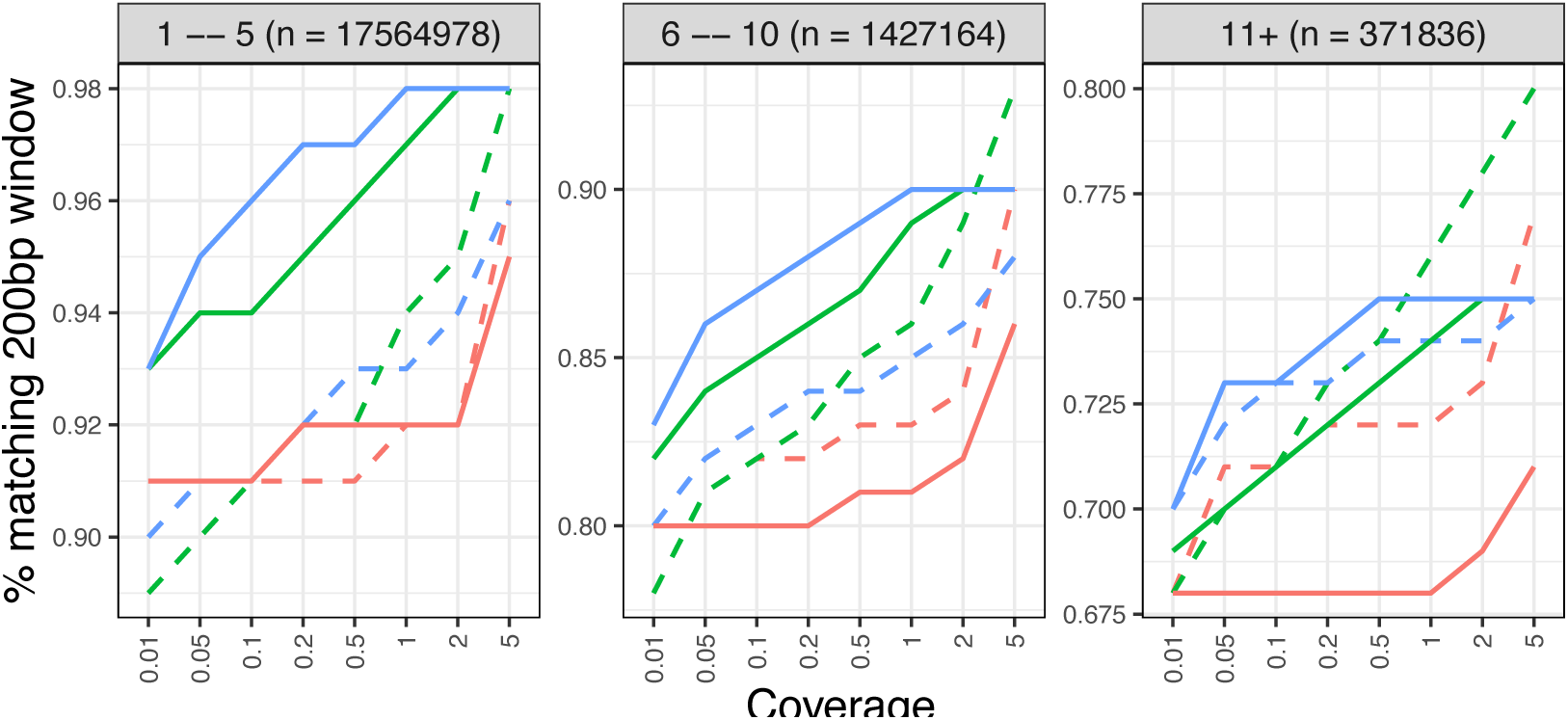
Window accuracy for diploid personalized genomes. The imputed sequence for each 200-bp windows anchored to a polymorphic site was compared to the ground-truth HG001 sequence. Results are stratified by the number of polymorphic sites in the window.

#### Computational performance

Figure 4 shows time and memory taken by these tool combinations on HG001, with similar observations noted for HG002. Glimpse used consistently less time and memory than

**Figure 4:**
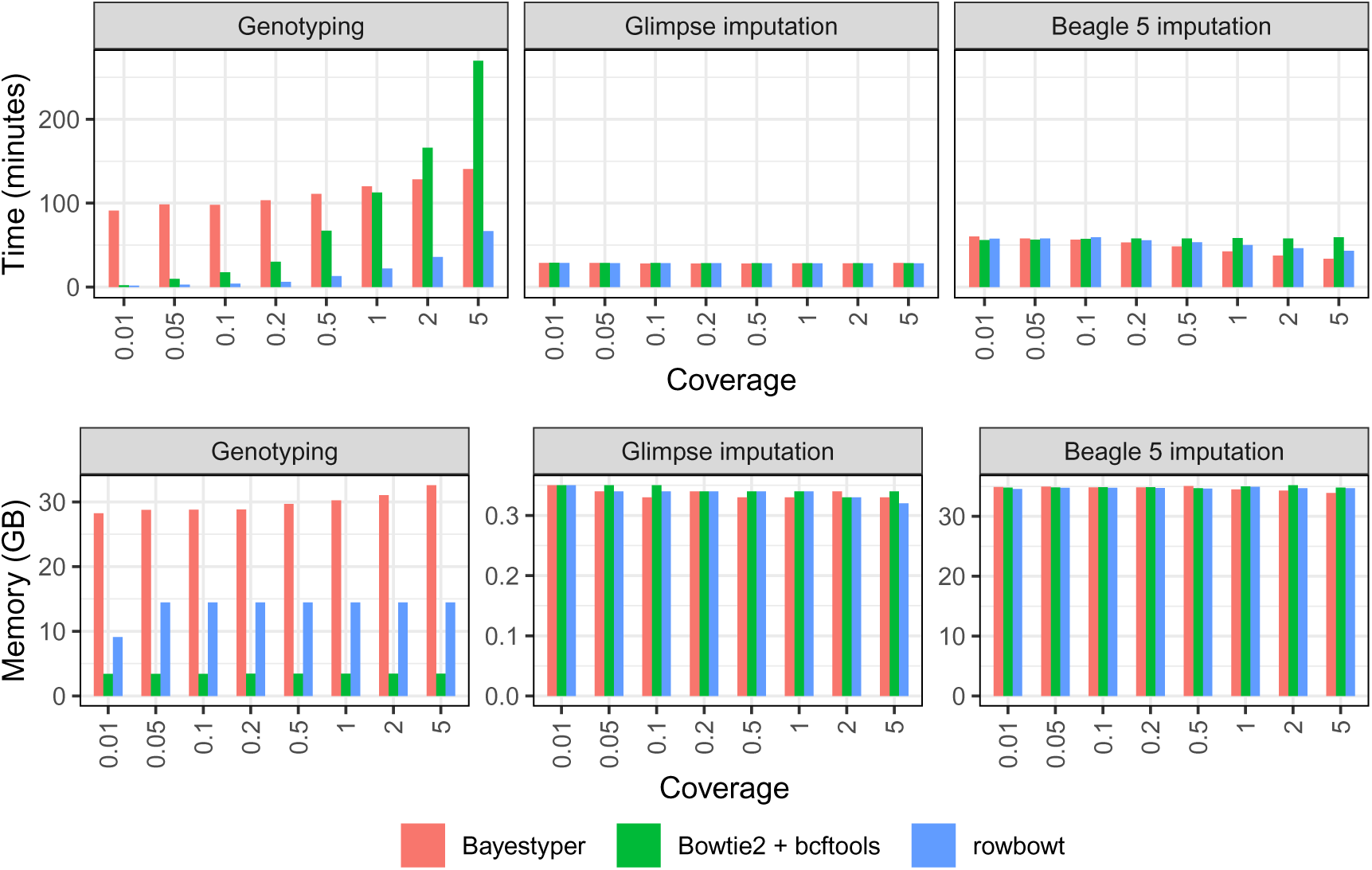
Computational overhead of genotyping (left) and imputation (middle: Glimpse, right: Beagle) in the personalization component of the impute-first workflow on HG001. In all cases, three genotyping methods are benchmarked. The Imputation methods are benchmarked three times, using the inputs from each of the 3 genotyping methods. Note the differences in scale for the plots in the bottom row.

Beagle. Specifically, Glimpse finished in about half the time used 1/100th as much memory as Beagle on average (Note the scale differences for the memory plots in Figure 4). Workflows using the Rowbowt genotyping tool were the fastest for the Genotyping step. For instance, at 1*×* coverage, the Rowbowt genotyping step took 22 minutes, whereas Bowtie2+bcftools took 112 minutes and BayesTyper took 120 minutes. Workflows using Bowtie2+bcftools were the most memory-efficient. At 1*×* coverage, Bowtie2+bcftools used 3.44 GB peak resident set size, while Rowbowt took 14.5 GB and BayesTyper took 30.2 GB.

#### Choices for Downstream Analysis

Given these results, for HG001, we selected the tool combinations rgc1, rgc5, bbgc5, and bbbc5. For HG002, we selected rgc5, bbgc5, and bbbc5. We chose these combinations both because of their high overall window accuracy, and because of their relatively high accuracy at lower coverage levels. We refer to these tool combinations by their short designators: rgc1, rgc5, bbgc5, and bbbc5, where these notations represent the provided distinct combinations of alignment and imputation methods.

### 2.3 Impact of personalized references on downstream results

Having produced personalized diploid references with the rgc1, rgc5, bbbc5, and bbgc5 personalization workflows, we next evaluated downstream results obtained using the personalized references. We compared the performance of downstream workflows using both personalized references as well as typical linear and graph-pangenome references. We used a combination of the BWA-MEM and VG Giraffe aligners, as seen in Table 2.

**Table 2:** Downstream workflows assessed.

| Aligner | Reference type | Used for |
| --- | --- | --- |
| BWA-MEM | `rgc1`-imputed diploid | Alignment score |
| BWA-MEM | Linear reference | Alignment score & variant calls |
| VG Giraffe | Linear reference<br>1000 Genomes phase-3 pangenome graph*<br>`rgc1`-imputed diploid<br>`rgc5`-imputed diploid<br>`bbgc5`-imputed diploid<br>`bbbc5`-imputed diploid | Allelic balance & variant calls |

\* For analyses using HG001, the 1000 Genomes phase-3 pangenome was modified to exclude HG001/NA12878 itself as well as family members NA12889 and NA12890.
*Note:* `rgc1`-imputed diploid is included in HG001 analysis.

#### 2.3.1 Alignment scores

We compared two sets of read alignments: one aligned to the the personalized diploid reference produced by the rgc1workflow, and another aligned to the linear GRCh38 reference (“GRC” for short). For each read that aligned to both, we asked whether its alignment score differed when aligned to the GRC versus to the personalized reference. 7.2% of the input reads had a different alignment score when aligned to one reference versus the other. Of these, 97% had higher alignment scores when aligned to the personalized reference. Among these reads, the alignment score improved by 7.7 points on average, equivalent in BWA-MEM’s scoring scheme to about 2 mismatches (each with a penalty of 4) or 1 small indel (with an open penalty of 6 and extension penalty of 1). Figure 5 shows all the alignment-score differences for reads having non-zero difference. These experiments show that by aligning to the personalized reference, we frequently remove penalties associated with the presence of non-reference alleles in the donor reads.

**Figure 5:**
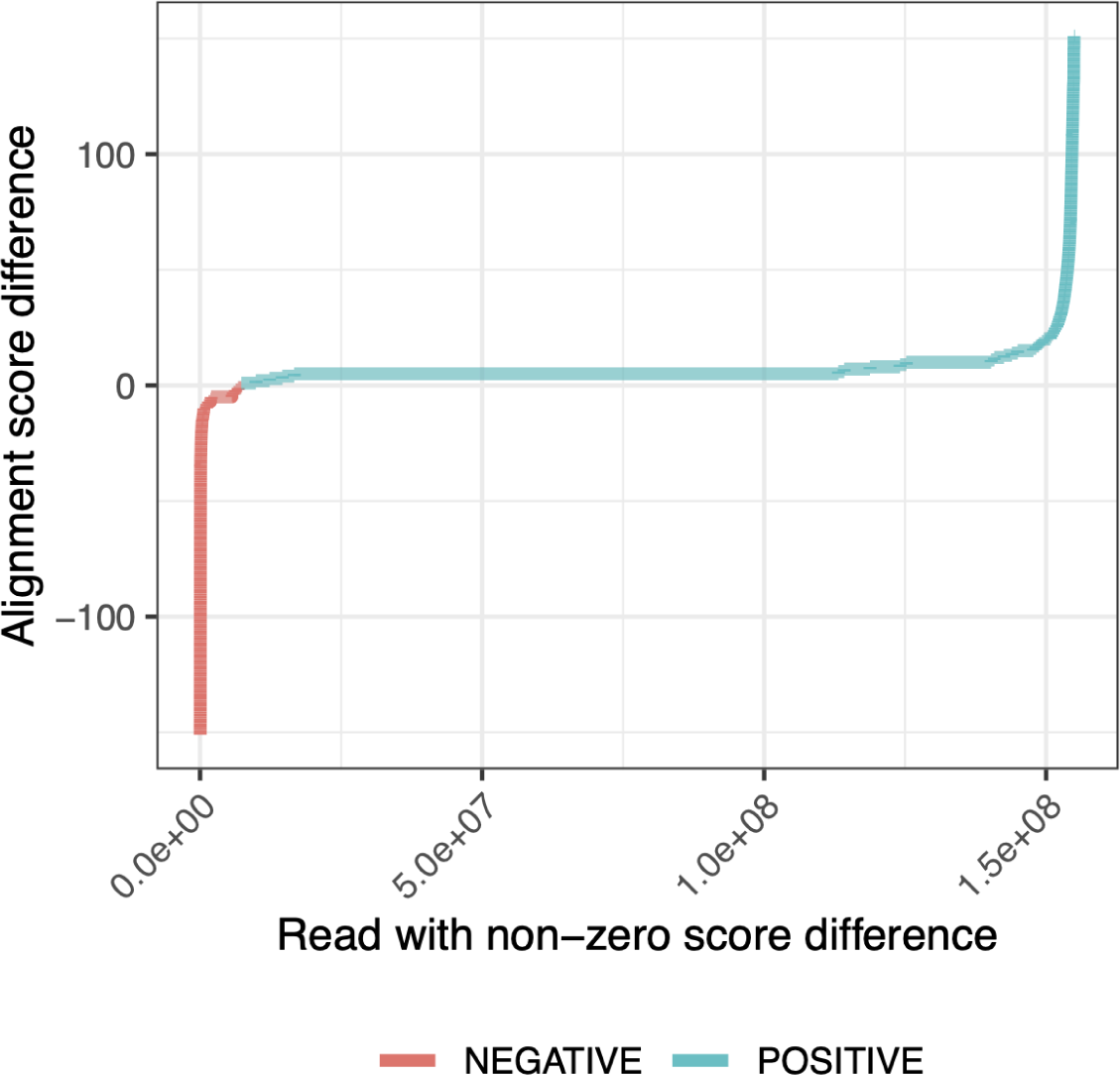
Difference in HG001 donor read alignment scores between personalized diploid reference (“Impute-first: rgc1”)and linear reference genome (GRCh38).

#### 2.3.2 Allelic balance at HETs

Next, we sought to measure how the personalized reference improves our ability to achieve a balanced representation of REF and ALT alleles at heterozygous (HET) sites in the donor. To do this, we began by using the Genome-in-a-Bottle (GIAB) HG001 v4.2 truth VCF file to identify sites where the NA12878/HG001 individual has a heterozygous varient. Following this identification, we used Biastools [40] to measure the allelic balance at each of these sites. Subsequently, we created an indel-balance plot, similar to those seen in previous studies of reference bias. Notably, we subsetted the analysis to focus on alignment intervals corresponding to the GIAB HG001 High-Confidence Regions (Figure 6A) and the Complex Medically Relevant Gene regions ([41, 42]) as depicted in Figure 6B. Figure 6 illustrates that alignments obtained using personalized references consistently achieve the most even allelic balance at HET sites. The personalized approaches show less bias than the pangenome reference, with the linear reference exhibiting the most imbalance. The trend is particularly evident at longer indels (left and right extremes), where impute-first references maintain a more even allelic balance compared to the others.

**Figure 6:**
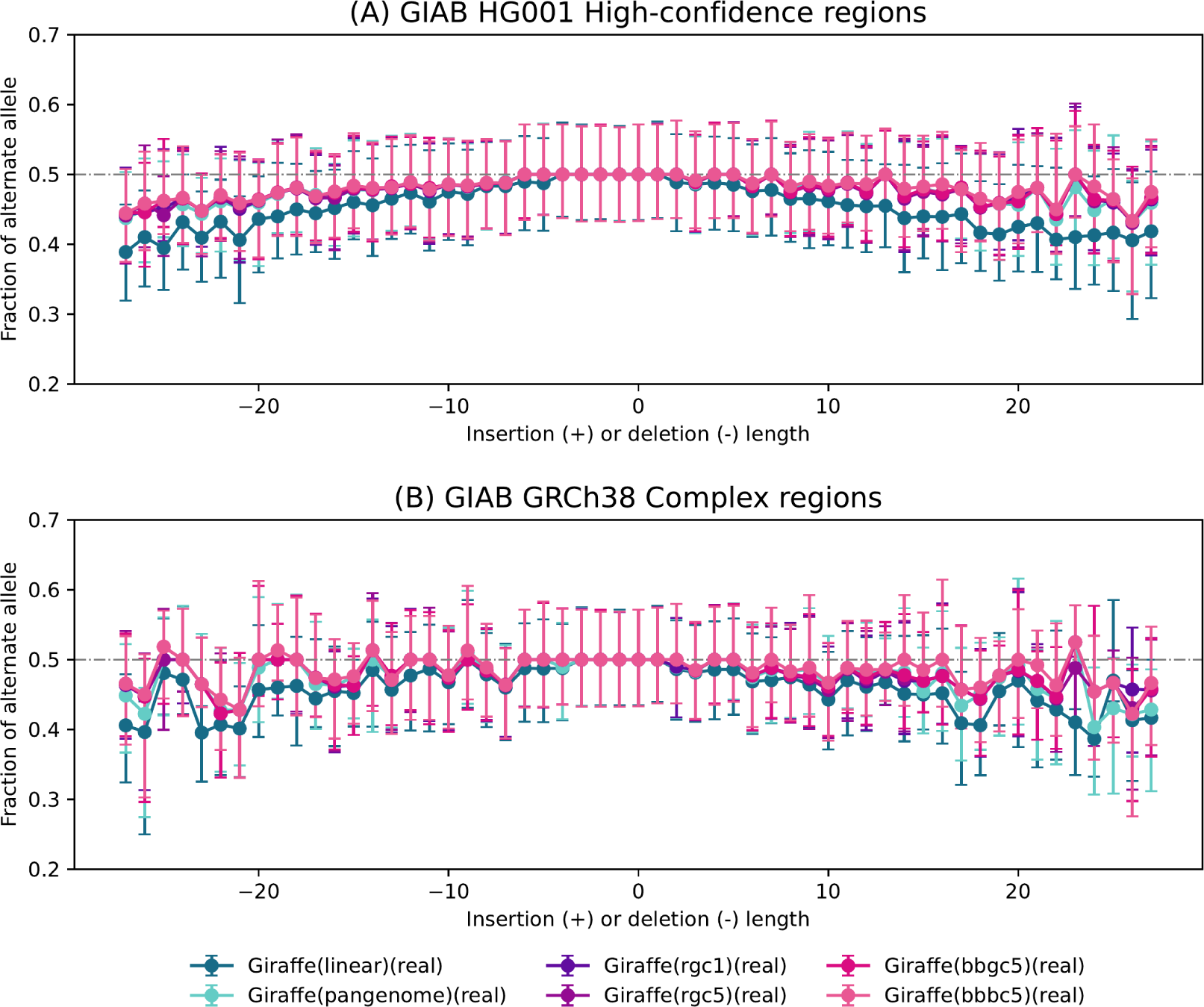
Bias-by-allele-length plots on HG001 donor reads generated for the 6 Giraffe-based downstream workflows. Variants are categorized by length: positive values for insertions, negative for deletions, and zero for SNVs at HET sites across the genome. (a) Allelic balance for indels and SNVs in GIAB HG001 high-confidence regions. (b) Allelic balance for indels and SNVs in GIAB GRCh38 Complex Medically Relevant Gene regions.

#### 2.3.3 Variant calling accuracy

Using RTG tools [37] and the GIAB truth VCF files for HG001 and HG002, we measured variant-level precision, recall, and F1 for both individuals. Measurements using Giraffe together with a linear or pangenome reference are labeled as “Giraffe (linear)” and “Giraffe (pangenome)” respectively. Measurements using impute-first workflows for HG001 are labeled with “Giraffe*” together with a designator (rgc1, rgc5, bbgc5, or bbbc5), and for HG002 with a designator (rgc5, rgc5, bbgc5, or bbbc5).

The impute-first workflows consistently achieved greater precision and recall than BWA-MEM as well as Giraffe when using its linear and pangenome indexes for both datasets (Figure 7). This was true both for SNVs and for indels (Figure 8). The results were consistent whether we measured variants only in high-confidence regions (Table S2 & Table S8), in the context of GRCh38 Complex Medically Relevant Gene (CMRG) regions (Table S3 & Table S11), or in the more challenging GIAB-labeled regions (Table S4 & Table S12). The results are also consistent across allele frequencies, with the rarest alleles exhibiting the largest recall gap between the impute-first workflows and the others (Table S7 & Table S14).

**Figure 7:**
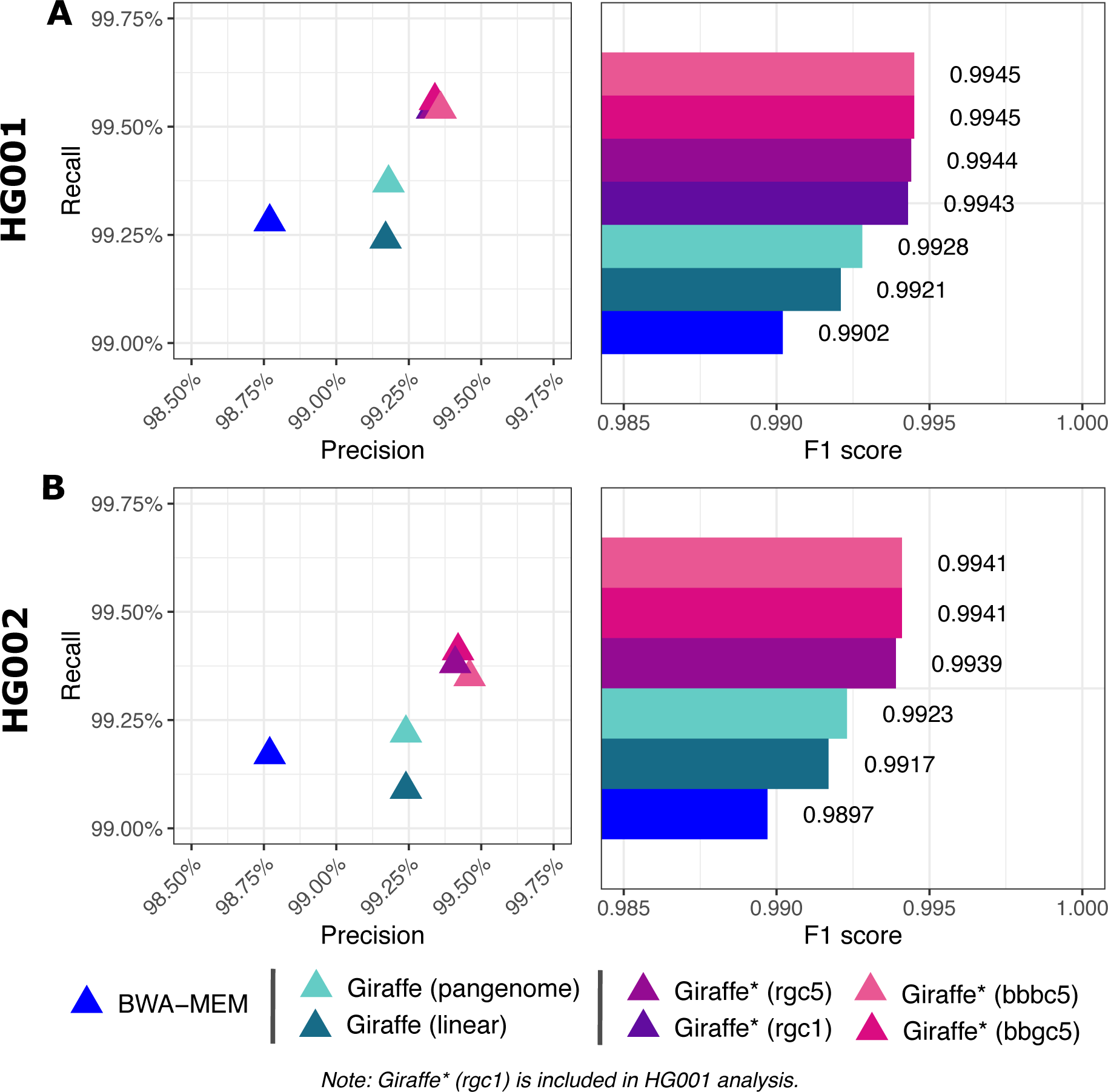
Precision-recall measures & F1 scores for overall variants. “Giraffe” indicates use of the standard VG Giraffe aligner. “Giraffe*” indicates use of VG Giraffe with a personalized reference produced by the personalization workflow mention in parentheses.

**Figure 8:**
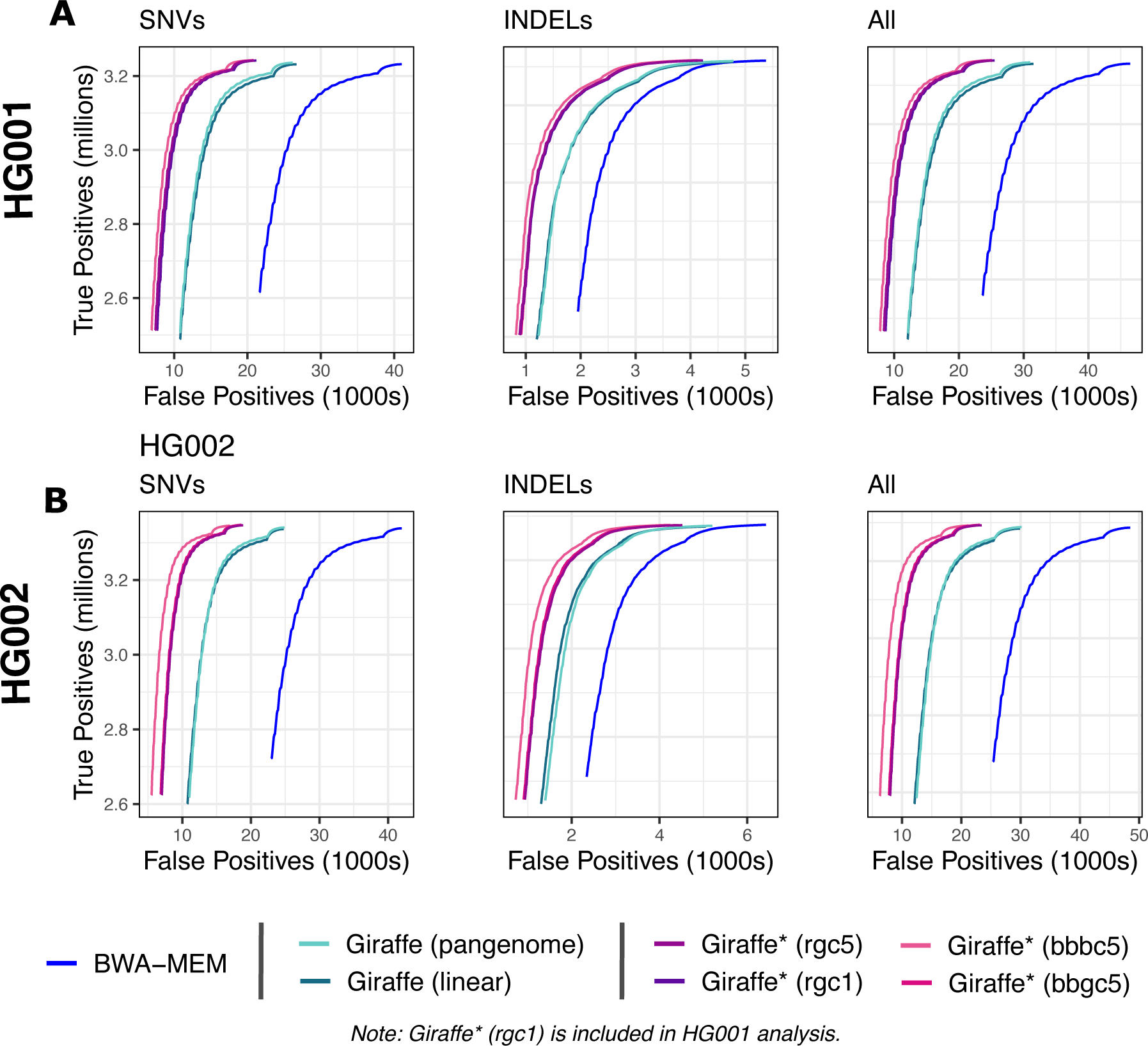
ROC curves from `vcfeval` ROC data files for SNVs, indels and overall variants. ROC curves are stratified by variant quality value threshold. “Giraffe*” indicates use of VG Giraffe with a personalized reference produced by the personalization workflow mention in parentheses.

The impute-first workflows did not differ much in their precision and recall across both datasets. However, the two workflows based on Bowtie2+bcftools performed slightly better than the two workflows based on Rowbowt. This difference was small compared to differences between the impute-first and non-impute-first workflows. There was no appreciable difference between the Glimpse-based versus the Beagle-based workflows.

For HG002, we extended our analysis using the T2T consortium’s new Q100 benchmark [43] [T2TQ100 HG002 v1.0 truth set and high-confidence regions]. The variant calling performance for HG002 demonstrated consistent improvements with impute-first workflows compared to BWA-MEM and Giraffe, both with linear and pangenome references (Figure S8, Tables S9 & S10). Within the GRCh38 Complex Medically Relevant Gene (CMRG) regions tailored for HG002 [44] [GIAB HG002 v1.0 CMRG truth & high-confidence regions], the impute-first workflows had slightly lower F1 score (0.9445–0.9524) compared to the Giraffe pangenome (0.9544), chiefly due to lower precision. When stratifying these CMRG results by allele frequency, we observed that the impute-first workflows performed particularly well for the rarest variants, and particularly when using Glimpse. (Table S15 & Table S16).

### 2.4 Computational efficiency

We also measured the computational overhead of all workflows, with particular attention to the overhead incurred by impute-first workflows. As noted, the impute-first workflow consists of distinct “personalization” and “downstream” phases. We evaluated the time and memory requirements for each of 6 workflows on HG001 (Figure 9). Two of the workflows were the Giraffe linear and Giraffe pangenome workflows, neither of which involved a personalization phase. The four others were the four impute-first workflows that each involved both personalization and downstream phases. For all 6 workflows, we measured the cost of building the reference index, though it should be noted that this step can be done once in an “offline” manner for the Giraffe linear and pangenome workflows, whereas it must be done in an “online” manner for the impute-first workflows.

**Figure 9:**
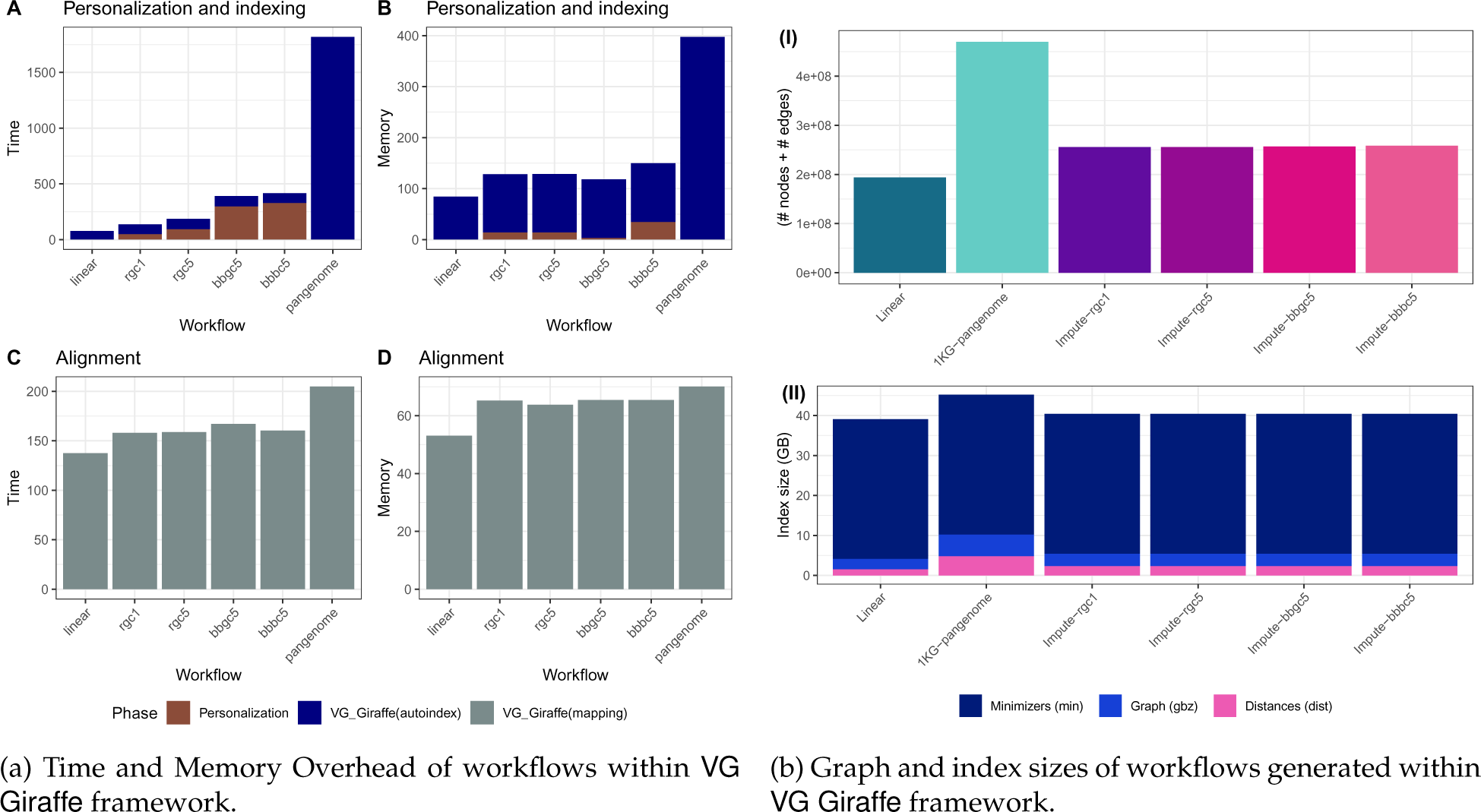
Computational efficiency of VG Giraffe-based workflows, considering time, memory footprint, and index size. Compared to typical pangenome VCF usage in VG Giraffe, the personalized VCFs used in Impute-first pipelines (*rgc1, rgc5, bbgc5, bbbc5*) offered more efficient reference representation and were also comparable to the linear method in terms of computational efficiency.

As shown in Figure 9a, building the Giraffe pangenome index took substantially more time than any of the other references. The impute-first references took about as much time to index as Giraffe’s linear reference, though the impute-first workflows required additional time for the other persoanlization steps, i.e. genotyping and imputation. The impute-first workflow required around 20*×* lower indexing time and 3*×* lower memory usage compared to the Giraffe pangenome workflow (Table S5). This benefit was also reflected in the size of the graph: the impute-first workflows yielded around half the number of nodes and edges compared to the Giraffe pangenome (Figure 9b & Table S6).

Figure 9 shows that, once the index has been built, the Giraffe pangenome workflow requires more time and memory than the other workflows, though these differences were smaller than the differences in indexing time and graph size. For example, the time required to align all the input reads to the Giraffe pangenome index was 27.8% greater than the time required to align to the impute-first bbbc5 reference.

Among the impute-first combinations, the Rowbowt combinations (especially rgc1) were most efficient, combining efficient genotyping in the personalization phase with lower downstream resource requirements versus the pangenome. Overall, the impute-first based personalized references demonstrated substantially improved downstream computational performance compared to using the pangenome reference within the VG Giraffe framework. Our analysis indicates that the impute-first approach can deliver accuracy improvements while maintaining reasonable computational overhead throughout the workflow.

We observed similar computational improvements on HG002 (Figure S7, with the rgc5-based workflow being the most efficient among the impute-first approaches.

A key caveat is that workflows using a personalized reference necessarily require the construction of a new reference index in order to leverage the personalized calls.

## 3 Discussion

We introduced a practical impute-first approach for genomic analysis with the goal of minimizing reference bias, even beyond what can be achieved with pangenome references. The workflow includes initial genotyping and genotype imputation steps, which produce a diploid personalized reference. This reference can be further leveraged by indexing it and using it as a basis for downstream alignment and variant calling.

The method yields improvements in all aspects measured: higher alignment scores, improved allelic balance at heterozygous sites, and enhanced downstream variant calls. VG Giraffe alignments using an impute-first personalized reference achieved greater variant-calling accuracy VG Giraffe with a pangenome reference. Further, the VG Giraffe index built on the personalized reference was more compact, reducing the computational burdens associated with building the index and aligning reads to it.

While past methods have explored the use of personalized references for avoiding reference bias, this study is the first to demonstrate a practical workflow that (a) performs personalization and downstream analysis with the same dataset, (b) integrates imputation tools like Glimpse and Beagle 5 that work with inputs derived from (possibly low-coverage) sequencing data, (c) outputs a diploid personalized reference genome for various downstream uses, and (d) works naturally together with a graph pangenome aligner to achieve both linear-genome-like computational overhead, and better-than-pangeome alignment accuracy.

We would note that while we concentrated on certain tools here, including BWA-MEM, VG Giraffe, Rowbowt and Glimpse, the framework we proposed is modular, and tools not assessed here could also be used. Our past work also assessed the PanGenie genotyper [45], for example, which could be integrated into the personalization phase [34].

In the future, it will be important to further improve the computational steps in the workflow’s personalization phase: genotyping and imputation. For instance, the recently described Minimal Positional Substring Cover (MPSC) framework [46] can solve a version of the imputation problem in time that does not depend on the number of haplotypes in the imputation panel. It is not immediately applicable, however, as it does not currently handle phased diploid imputation.

Another avenue for improvement may be to combine results from multiple imputation tools to improve personalization accuracy. Such benefits were reported previously in ancestral DNA analysis studies [7].

While our method shows promising results with whole-genome DNA sequencing data, more work is required to establish its applicability to other assays like exome sequencing and RNA-seq. These assays exhibit uneven coverage across the genome, which complicates the initial genotyping and imputation steps. For exome sequencing, coverage is deep in exonic regions and shallow elsewhere, leading to higher genotyping and imputation accuracy in exons; accuracy will then presumably fall off in areas flanking the exons, and only low accuracy will be achieved between exons. RNA-seq poses an additional problem duew to variability in expression levels also complicates genotyping, since lowly-expressed genes may not provide sufficient data for genotyping and imputation.

We opted to use the HGSVC2 reference panel in the personalization phase of our workflow because that panel was compiled using advanced long-read assembly methods that allowed it to survey complex structural variants. However, more work is required to determine which are the most important features the imputation panel should have to minimize reference bias and maximize downstream accuracy. While we opted for a panel with relatively few individuals but a relatively wider survey of structural variants, it could also be advantageous to choose a panel with more individuals but a narrower range of SVs, such as that produced by the 1000 Genomes project. Notably, the recent Glimpse2 algorithm [47] is able to handle BioBank-scale reference panels, as demonstrated using the 150K samples in the UK BioBank.

We used the GATK HaplotypeCaller to call variants here. We did not use DeepVariant, because its internal model is already “trained” to undo the effects of reference bias, which it can readily observe in the training data that it obtains using alignment tools that produce typical levels of reference bias. More study is needed to find how the most effective tools for avoiding bias “upstream” can be combined with the most effective bias-avoiding tools “downstream.” This requires tighter feedback between the two, e.g. with downstream models being trained on alignments produced by the best bias-avoiding upstream tools.

## 4 Methods

Figure 1 illustrates the impute-first workflow. First, a set of reads is sampled from the full set of input reads. Next, the sampled read set is used to obtain rough genotypes for the donor. The rough genotypes are passed to an imputation tool, which imputes and phases both donor haplotypes, yielding a personalized diploid reference genome. This is the “personalization” portion of the workflow, depicted in Figure 1A.

In scenarios where high-quality variant calls are desired, the workflow can be continued further downstream, as depicted in Figure 1B. In this case, the diploid personalized reference will be indexed and used to analyze the full input read set.

### 4.1 Personalization

#### 4.1.1 Sampling

To minimize the computational burden of personalization phase, we tested its accuracy and efficiency when run on only a sample of the input read set. For instance, if the input reads total 30*×* average coverage, the personalization workflow might use a random subset of those reads totaling 1*×* average coverage. Past work has shown that genotype imputation can be performed with excellent accuracy even when the input reads represent a low level of average coverage [38, 48].

#### 4.1.2 Genotyping

The next step obtains a set of “rough” variant calls based on the subsampled reads. Since the sampled input might comprise a low level of average coverage, the genotypes obtained in this step are not expected to be high quality. This is acceptable in practice because the imputation step that follows combines this rough genotyping information with the large database of high-quality phased genotypes available in the imputation panel to produce an output that is much higher quality.

This step of the personalization workflow is modular; various tools could be inserted here to accomplish the rough genotyping task. In our experiments, we used four different specific toolsets, described in the following subsections.

##### Rowbowt

Rowbowt [34] uses an *r*-index [13] together with an auxiliary *marker array* structure to achieve a compressed pangenome index where genotype evidence is easily tallied. First, rowbowt takes a Variant Call Format (VCF) file and a corresponding FASTA file as input, where the FASTA provides the reference sequence and the VCF describes the variants present on the haplotypes to be indexed. It then indexing the index using the prefix-free parsing algorithm [49].

We used Rowbowt to index the HGSVC2 reference panel, which comprises 34 samples, or a total of 68 haplotypes. Since some of our experiments analyzed reads from the individual with accession NA12878, we first excluded NA12878’s two haplotypes before building the Rowbowt index. In addition to the HGSVC2 haplotypes, we also included the GRCh38 primary assembly as one of the indexed references. The marker array’s window size *w* was set to 19, the value used and tested in the Rowbowt study [34]. Rowbowt computes and reports genotype likelihoods (following a method similar to past ones [50]), which can then be provided to the downstream imputation tool.

##### Bowtie2 + bcftools

The genotyping procedure against the linear reference genome involved a series of steps. Initially, raw reads were aligned to the human reference genome, GRCh38 using Bowtie 2 v2.4.2 (Bowtie2) [31] with default parameters and 16 threads. After alignment, a pile-up of overlapping reads was generated for each genomic position using BCFtools v1.13 (`bcftools`), [51] with `bcftools mpileup` command using default settings. This pile-up information was then used for variant calling, accomplished through the `bcftools call` command with a multi-allelic calling model, retaining default parameters.

To ensure the accuracy and reliability of identified variants, the low-quality attributes were filtered out by excluding VCF records with INFO/QUAL values below 20 and INFO/DP values exceeding 100. It’s important to note that both pile-up generation and variant calling stages were conducted using single-thread configurations due to the lack of multi-threading support in these respective analysis modules used.

##### BayesTyper

BayesTyper [33] operates by matching k-mers from input reads to k-mers stored in a de Bruijn graph that represents known polymorphisms and their neighboring contexts. This evidence is then leveraged to perform genotype calls through a generative model. The genotyping process involves constructing variant graphs using alleles defined by a predetermined set of variants in a VCF file, followed by read-to-graph mapping by aligning read k-mers to graph nodes.

In our experiments, we have used BayesTyper v1.5 (BayesTyper) and built a compatible VCF file to contain all relevant variant set from the HGSVC2 reference panel. For enhanced computational efficiency, 32 threads are employed during k-mer counting and bloom filter construction, and 16 threads during the genotyping phase.

### 4.2 Diploid genotype imputation

#### 4.2.1 Beagle

We used Beagle v5.1 (Beagle) [17] to phase and impute genotypes based on genotype inputs taken from the four genotyping methods. We executed Beagle with default parameters but using 32 simultaneous threads. We performed phasing and imputation on a per-chromosome basis. For the imputation reference panel, we used the 34 samples from the HGSVC2 project, excluding NA12878 and family members (NA12889 & NA12890). The genetic map files were sourced from the Beagle web resource [52].

#### 4.2.2 GLIMPSE

We used Glimpse v1.0.0 [18] to impute genotypes based on genotype inputs taken from the four genotyping methods. We used the same reference panel and genetic map as for the Beagle experiments, first converting the genetic map to Glimpse’s format. We executed Glimpse with 32 simultaneous threads. We ran all of Glimpse’s distinct modules: `chunk`, `ligate`, `phase`, and `sample` with default parameters. As we did for Beagle, we conducted the imputation on a per-chromosome basis.

### 4.3 Construction of personalized reference and index

#### 4.3.1 Personalized linear reference

We used `bcftools consensus` to construct a diploid reference genomes in the FASTA format. We used `bcftools consensus` to insert the imputed, phased variants from the rgc1-personalization workflow into the GRCh38 reference, creating two personalized haplotypes. We next created indexes for these haplo-types (`Haplotype 1`, `Haplotype 2`) as well as for the GRCh38 reference using BWA-MEM’s (`bwa index`) command with default parameters.

#### 4.3.2 Personalized graph reference

We generated personalized reference graphs using `vg autoindex --workflow giraffe`, using `vg` version 1.46.0 for HG001 and a later `vg` version 1.55.0 for HG002. We used default parameters and 32 simultaneous threads for both. As inputs, we provided both the GRCh38 reference FASTA and the personalized diploid variant calls from the corresponding personalization workflows for each individual. The resulting graphs were “personalized reference graphs” HG001’s and HG002’s unique variants respectively. That is, for variants that were called homozygous ALT compared to the GRCh38 reference, those positions were switched to the ALT allele in each respective graph. For variants that were heterozygous, the graphs will contain alternate paths for both of the two alleles. To assess the size and complexity of these graphs (e.g. for Figures 9b & S7b), we used the `vg stats` command to obtain the size of the graph. Additionally, for further downstream evaluations and comparisons that include index sizes and indexing performance, we performed index constructions for linear (without VCF) and pangenome (1kGP phase 3 VCF) combinations using the GRCh38 reference fasta in the `autoindex` workflow with `giraffe` for both individuals.

### 4.4 Alignment and variant calling

#### 4.4.1 Alignment with linear aligner

For the “alignment scores” analysis discussed in Figure 5, we used BWA-MEM with default parameters to align the full set of HG001 donor reads to the standard GRCh38 reference. We also used BWA-MEM to align all the reads to the both haplotypes of the personalized diploid reference produced using the “impute-first: rgc1” method. For the personalized reference, reads were aligned separately to both of the personalized haplotypes. After this, the alignment with the maximum scores between the two haplotypes was chosen for the analysis.

#### 4.4.2 Alignment with VG Giraffe

Using the personalized reference graph indexes created through the VG Giraffe `autoindex` workflow, we employed the VG Giraffe mapping module (`vg giraffe`) to align the full set of HG001 donor reads. For improved accuracy, we supplied read-specific statistics to the algorithm, including the fragment length mean and standard deviation as inferred for the HG001 dataset by BWA-MEM. This alignment process resulted in standard BAM files for the respective combinations, serving as input for subsequent analyses. We followed the same alignment procedure for all the Giraffe-based workflows. For HG002, the alignment was performed using a later release of `vg` version (v1.55.0), which infers read-specific statistics by default, unlike the earlier `vg` version (v1.46.0) used for HG001.

#### 4.4.3 Variant calling with GATK HaplotypeCaller

In the variant calling phase, BAM alignment outputs from VG Giraffe workflows (linear, pangenome, impute-first combinations) from both HG001 and HG002 were processed. After formatting the BAMs to GATK (v4.2.6.1) standards, GATK HaplotypeCaller (v2.24.1) [53] was applied to each BAM with the GRCh38 genome as the reference. The `HaplotypeCaller` identified potential variants for both HG001 and HG002 from the corresponding BAMs. These results were captured in VCF files and benchmarked against the GIAB truths for HG001 and HG002, respectively.

### 4.5 Evaluation

#### 4.5.1 Measuring genotyping and imputation accuracy

We analyzed the accuracy of the personalization workflows’ diploid genotypes in two ways. First we considered allele-by-allele precision and recall, considering the alternate (ALT) allele calls from the sample-specific truth VCF to be the positive class. Specifically, every diploid genotype called by a method is considered as a pair of individual allele calls. If a given allele call is an alternate (ALT) allele and there is at least one ALT allele present in the true diploid genotype at that site, it counts as a true positive (TP). If the given allele is a reference allele (REF) and there is at least one REF allele in the true diploid genotype, this is a true negative (TN). If the given allele is an ALT but the true genotype is homozygous REF, this is counted as one false positive (FP). Finally, if the given allele is REF but the true genotype is homozygous ALT, this is counted as one false negative (FN).

Second, we considered precision and recall with respect to sites that were either truly heterozygous or called heterozygous. If a heterozygous call made by a method is truly heterozygous, this was counted as a true positive (TP). False positives, false negatives, and true negatives are defined accordingly.

In both cases, precision and recall are computed as:

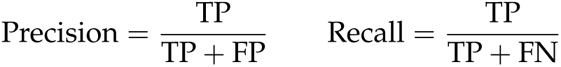

#### 4.5.2 Measuring window accuracy

To evaluate the accuracy of the short-range phasing surrounding each marker, for both haplotypes in the truth set, we extracted the 200bp flanking sequences around each polymorphic site and tallied their presence/absence in the test/impute-first genotypes. We call this “window accuracy” and calculate it as:

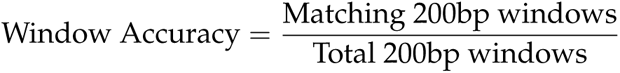

#### 4.5.3 Measuring accuracy of personalized genome alignments

For the analysis presented in Results 2.3 and Figure 5, reads aligned with BWA-MEM were selected based on variations in alignment scores across the GRCh38 and the two haplotype copies constructed from the HG001 “impute-first: rgc1” variants. Emphasis was placed on reads with non-zero score differences between the references. This subset of reads was then analyzed for both positive and negative score differences. For each read, the optimal alignment score from either of the two haplotype copies was determined and considered as the personalized score for that read. Following this, a comparison was conducted to assess the differences between the alignment scores of the GRCh38 and the personalized reference.

#### 4.5.4 Measuring allelic balance at HETs

To quantify reference bias, we ran *biastools* including its context-aware assignment algorithm [40]. This tool examines the alignment evidence overlapping heterozygous variants. We ran *biastools* separately on per-chromosome HG001 SAM files, extracted from the workflow-generated BAM outputs. At heterozygous sites, *biastools* tallies and reports the allelic balance, i.e. the number of ALT-carrying reads overlapping the site divided by the sum of the ALT- and REF-carrying reads overlapping the site. To identify heterozygous sites, *biastools* was given the GIAB HG001 v4.2.1 truth VCF. We configured *biastools* to limit its analysis to HETs contained in the GIAB-designated high-confidence regions of HG001 (Figure 6a). We repeated this, asking *biastools* to limit its analysis to HETs in the GRCh38 challenging, medically relevant genes (CMRGs) (Figure 6b). Figure 6 first partitions HET sites according to the length of the inserted or deleted sequence, with SNVs considered to be length zero. For each length category, the plot shows median allelic balance.

#### 4.5.5 Assessing variant call accuracy

We ran `rtgtools vcfeval` using the latest HG001 and HG002 v4.2.1 truth VCFs against each of the callsets generated from GATK on feeding corresponding BAMs of linear, pangenome, impute-first workflows generated within VG Giraffe. The evaluations were performed in aggregate (overall variants) mode. For the high-confidence regions, we also stratified by SNVs versus indels.

We used the RTG Tools [37] `vcfeval` command (v3.12.1) to evaluate variant calling accuracy. Specifically, we compared variant calls made by the alternate workflows to the true genotypes reported in the GIAB HG001 and HG002 v4.2.1 truth VCFs [36] respectively. For Figures 7 & 8 and Tables S2 & S8, we limited the analysis to variants in regions called “high confidence” in HG001 by GIAB. For Tables S3,S11 and S13, we limited the analysis to the challenging, medically relevant genes (CMRGs) discussed by Wagner et al. [44]. For Tables S4 and S12, we reported results for a few other challenging subsets of genomic regions, specifically the “MHC,” “alldifficultregions,” “allOtherDifficultregions,” and “allowma-pandsegdupregions” regions as defined by GIAB.

We used the default optimized threshold parameters in `rtgtools vcfeval` to summarize standard precision, recall, and F1 accuracy metrics for each callset compared to the GIAB truth. We used the ROC output from `vcfeval` to assess performance (e.g. in Figure 8) on the high-confidence regions for SNP, indel, and overall variants.

## 6 Availability of data and materials

- GRCh38 reference fasta
- HG001 Fastq files: SRR622457, HG001.novaseq.pcr-free.30x.R1.fastq.gz, HG001.novaseq.pcr-free.30x.R2.fastq.gz
- HG002 Fastq files: HG002.novaseq.pcr-free.30x.R1.fastq.gz, HG002.novaseq.pcr-free.30x.R2.fastq.gz
- HGSVC, phase2 haplotype reference panel
- Genetic maps
- GRCh38 phase3 1kGP callset
- GIAB HG001 v4.2.1 truth VCF & High-confidence regions
- GIAB HG002 v4.2.1 truth VCF & High-confidence regions
- GRCh38 HG2-T2TQ100-V1 (T2TQ100 HG002 v1.0) truth VCF & High confidence regions
- GIAB GRCh38 v3.1 Genome stratifications
- GIAB GRCh38 Complex Medically Relevant Genes (CMRG) regions
- GIAB HG002 v1.0 CMRG truth VCF & High-confidence regions

Scripts for the experiments described in the article are at Impute-First Alignment framework.

## 7 Authors’ contributions

KV, TM and BL designed the method. KV wrote the software and performed the experiments. KV and BL wrote the manuscript. All authors read and approved the final manuscript.

## Supplementary Figures

**Figure S1:**
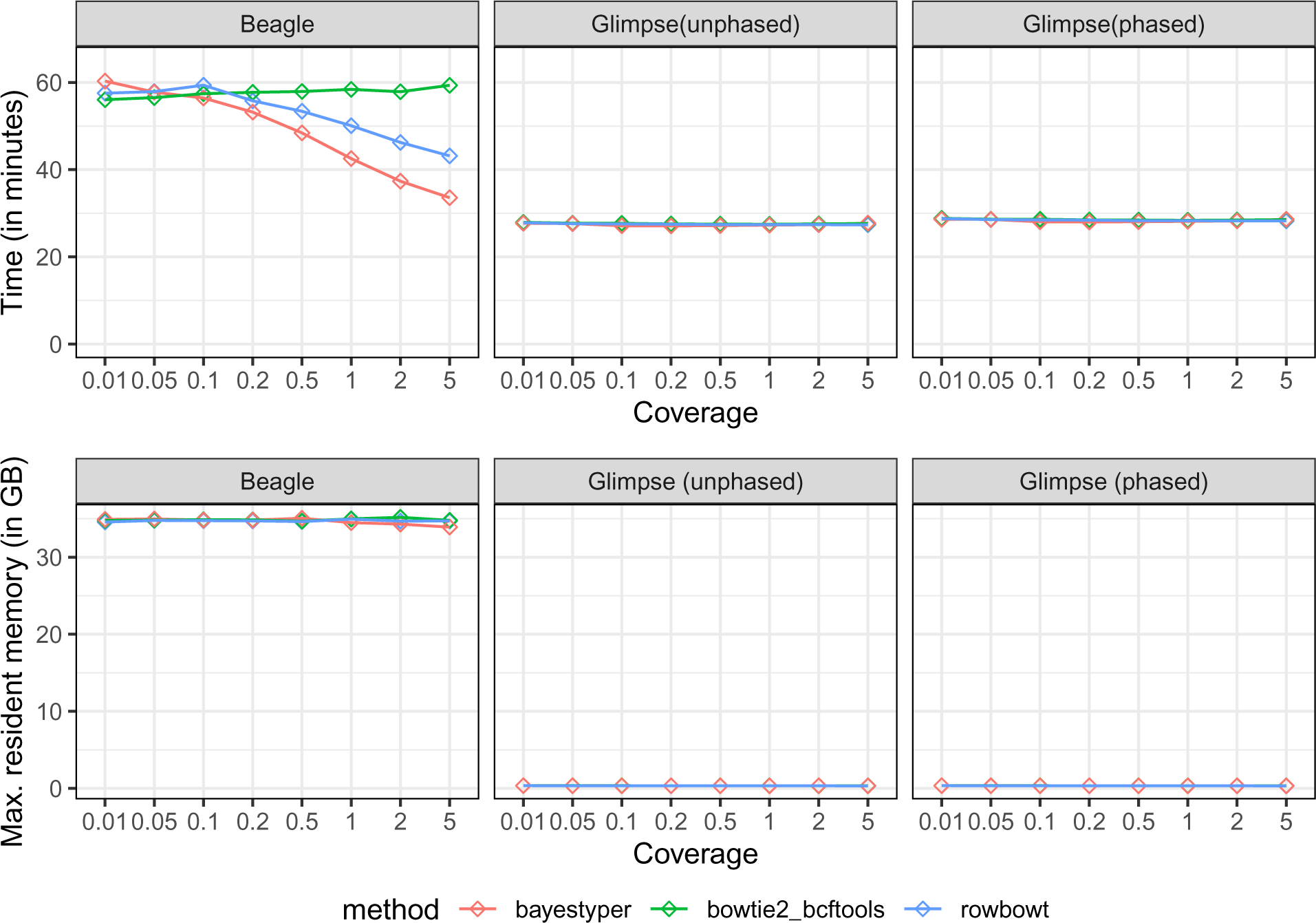
Computational overhead for Beagle and Glimpse stratified by the genotyping tool used upstraem, and as a function of the read coverage provided to the genotyper. Measurements for Glimpse were taken in both its phased and unphased modes.

**Figure S2:**
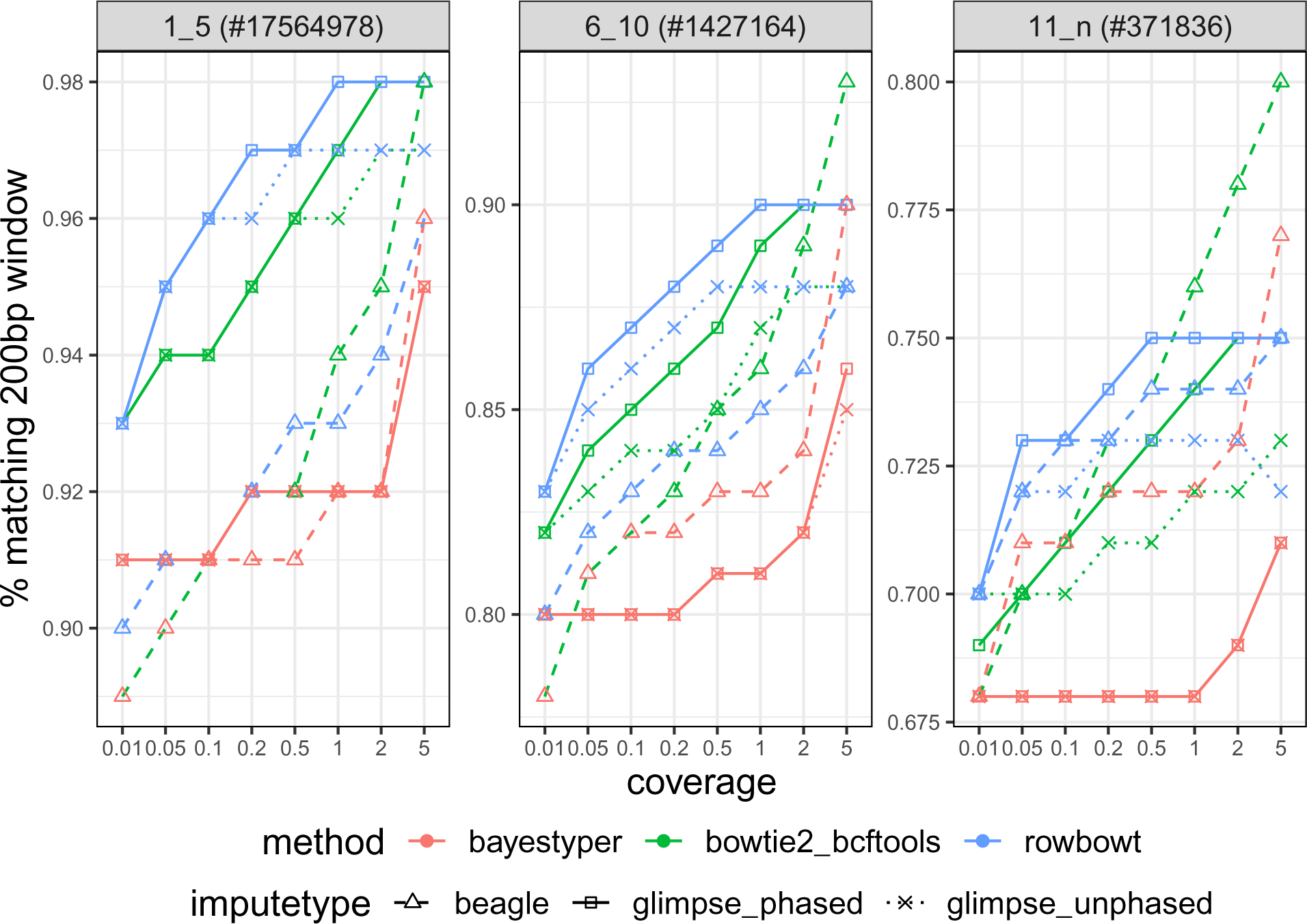
Window Accuracy for Beagle and Glimpse (in both phased and unphased modes) for different methods on diverse coverages.

**Figure S3:**
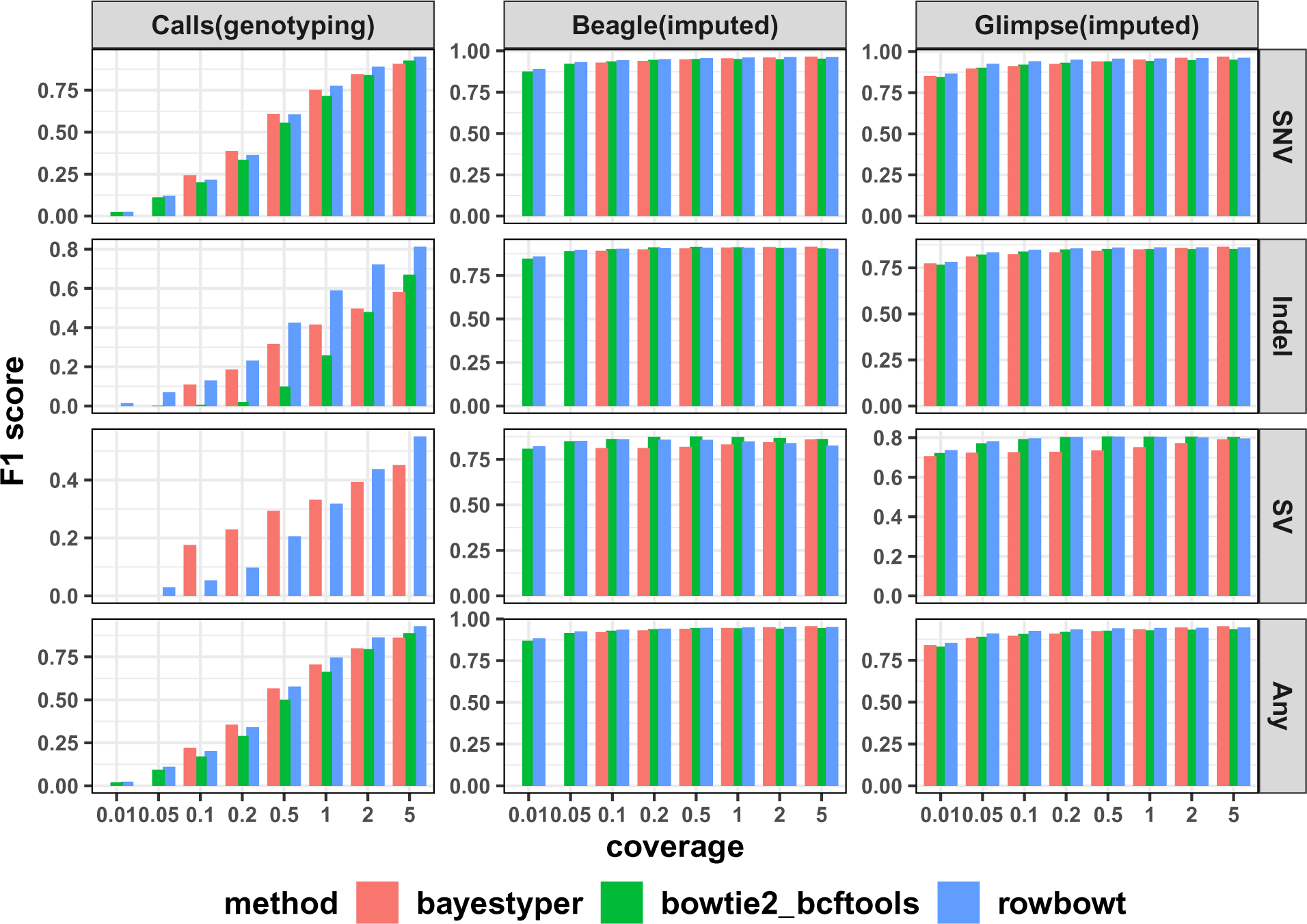
F1 score of alternate allele calls (ALT), stratified by variant type, for the personalized genomes constructed using each alignment/genotyping method in the Impute-first alignment workflow.

**Figure S4:**
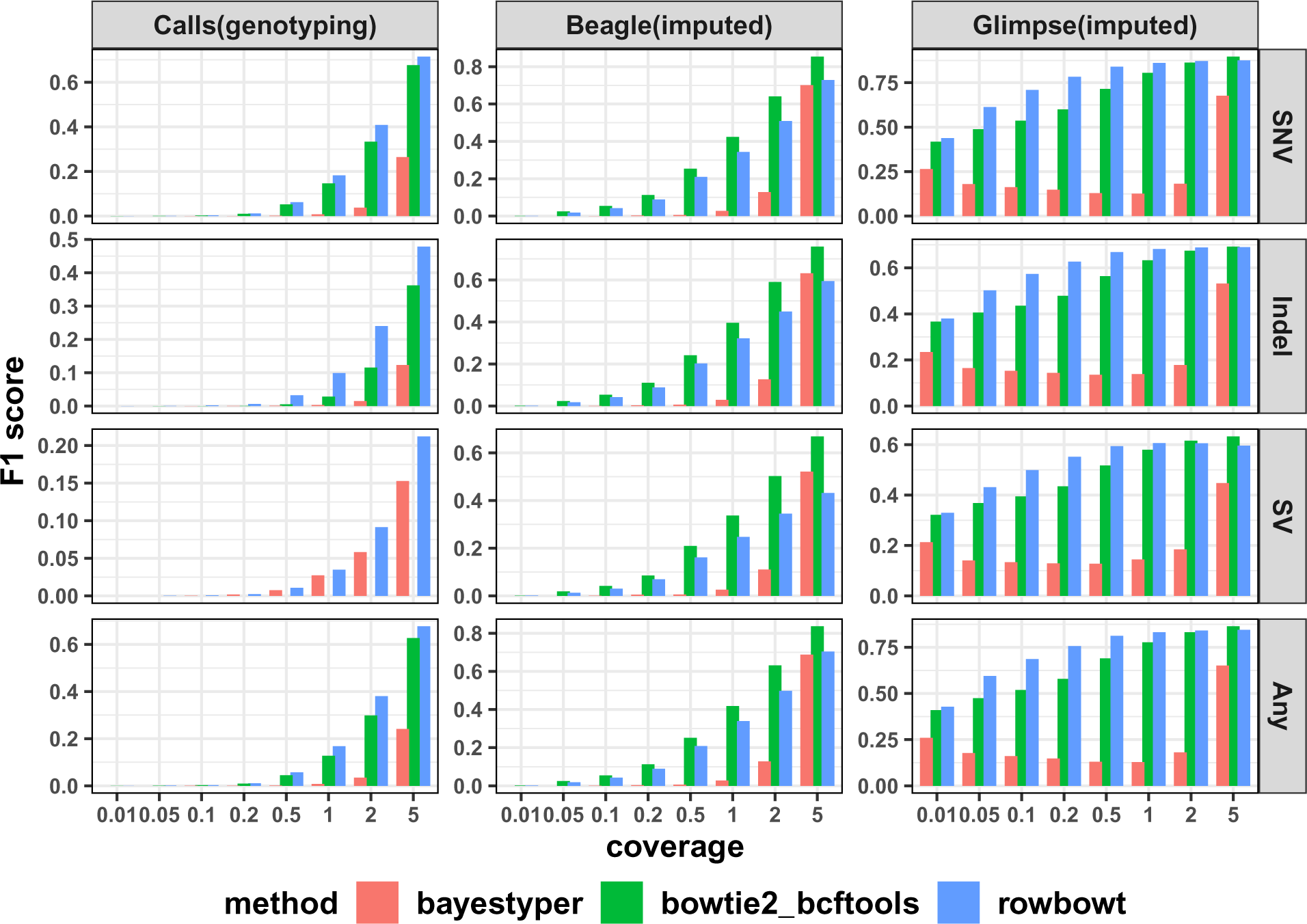
F1 score of heterozygous calls (HET), stratified by variant type, for the personalized genomes constructed using each alignment/genotyping method in the Impute-first alignment workflow.

**Figure S5:**
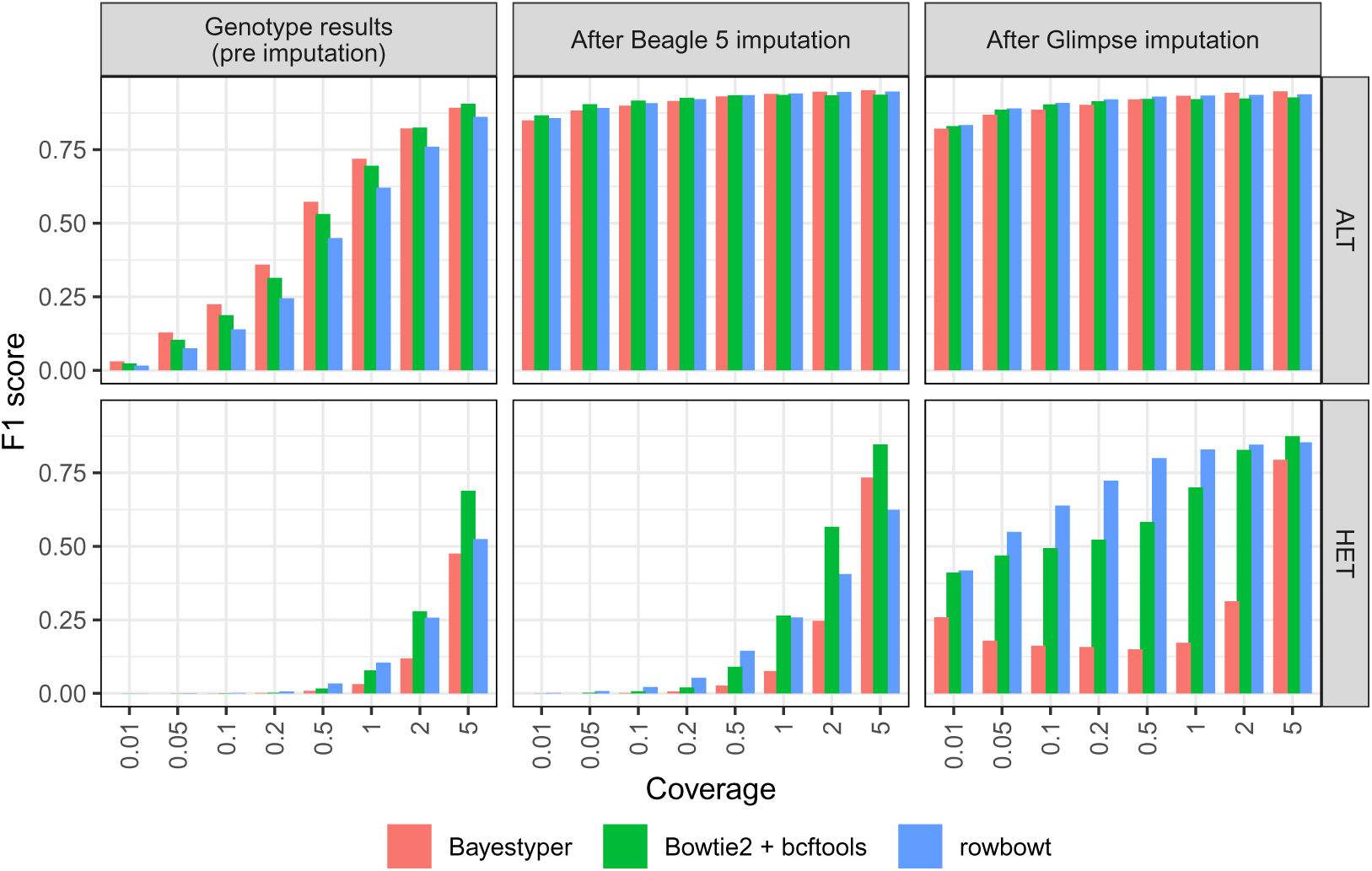
Aggregate F1 scores of alternate allele calls (ALT) and heterozygous calls (HET) across all variant types, generated using each alignment/genotyping method in the Impute-first alignment workflow on HG002.

**Figure S6:**
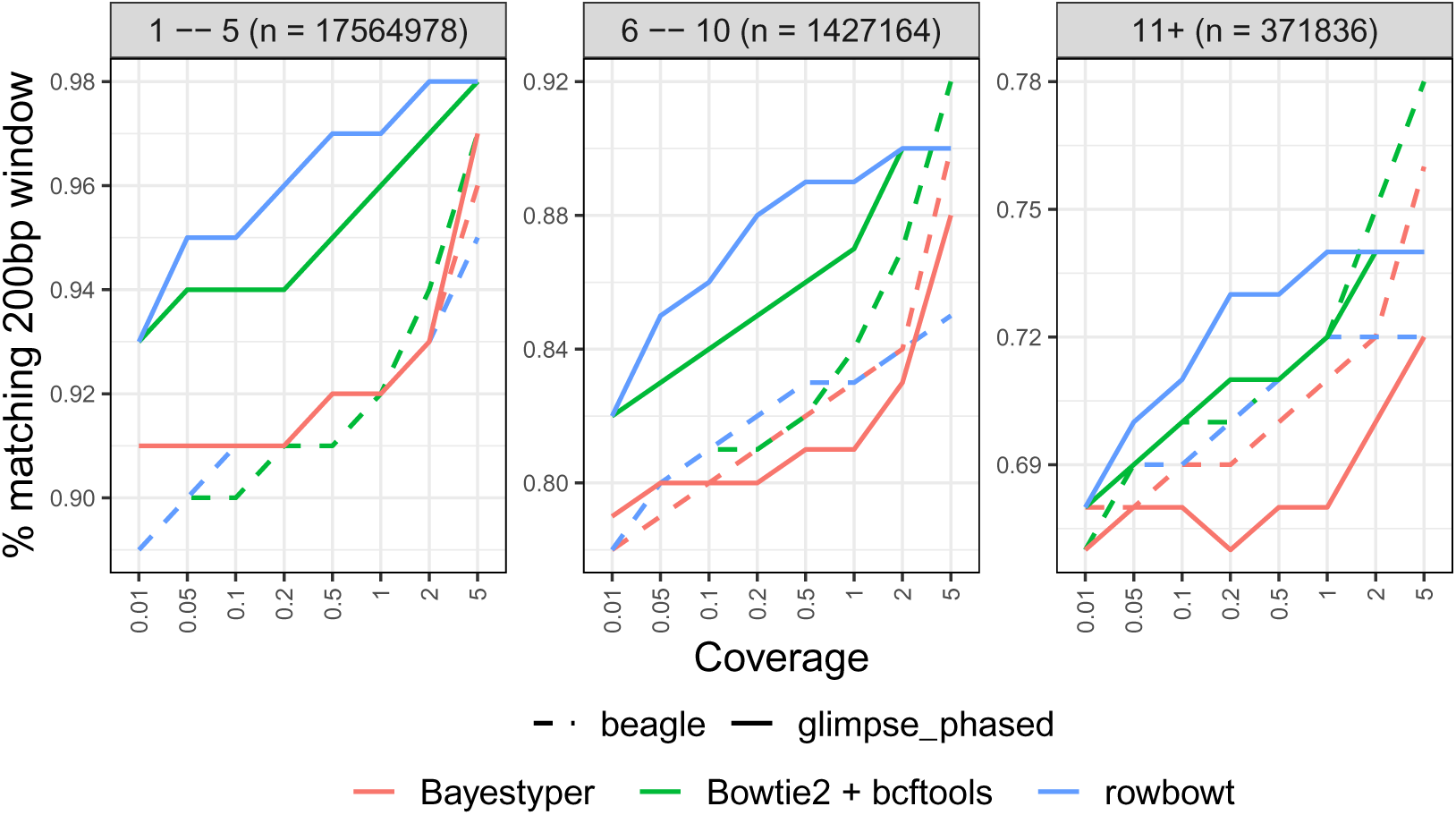
Window accuracy for diploid personalized genomes using HG002 data. The imputed sequence for each 200-bp windows anchored to a polymorphic site was compared to truth NA24385 sequence. Results are stratified by the number of polymorphic sites in the window.

**Figure S7:**
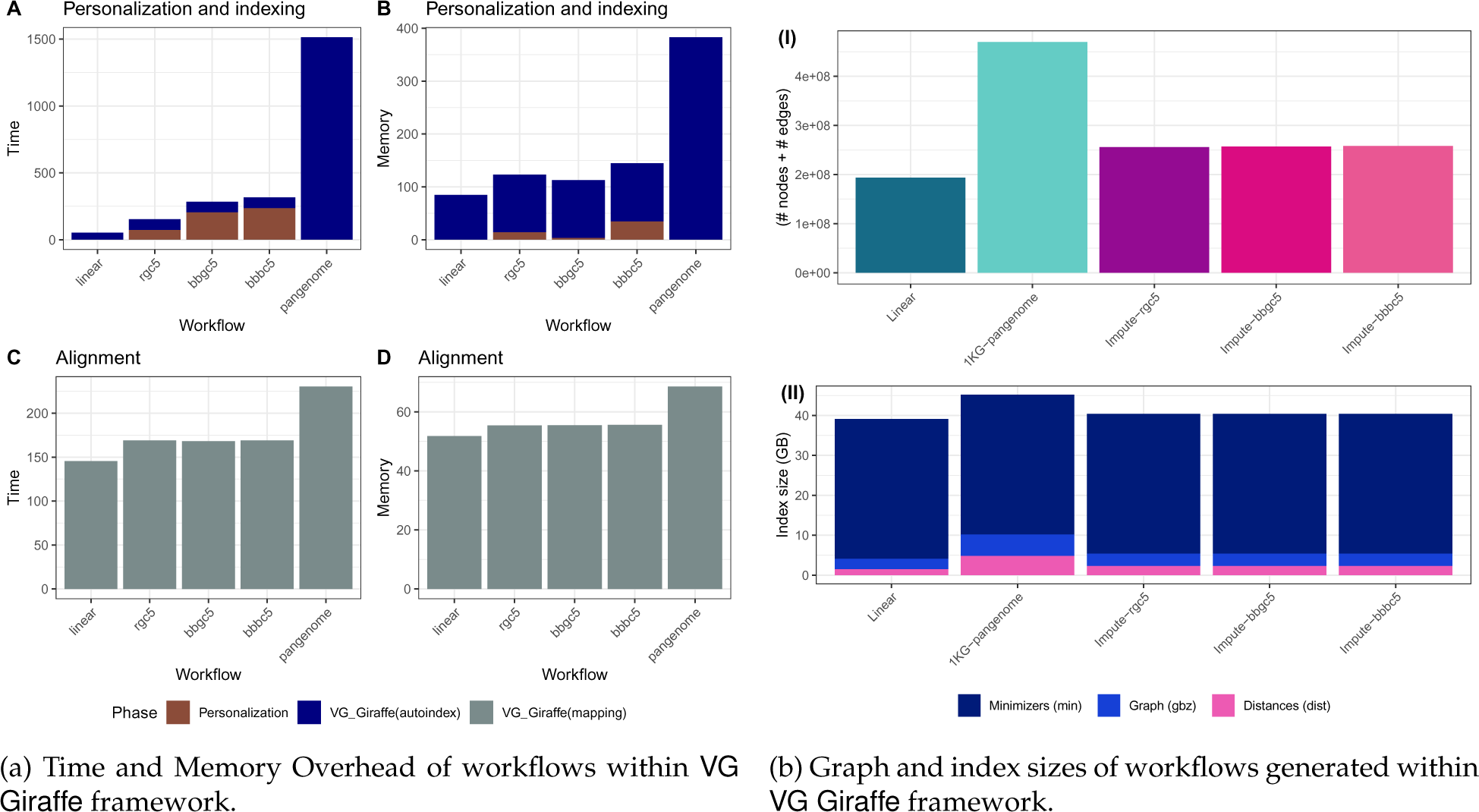
Computational efficiency of VG Giraffe-based workflows on HG002, considering time, memory footprint, and index size. Compared to typical pangenome VCF usage in VG Giraffe, the personalized VCFs used in Impute-first pipelines (*rgc5, bbgc5, bbbc5*) offered more efficient reference representation and were also comparable to the linear method in terms of computational efficiency.

**Figure S8:**
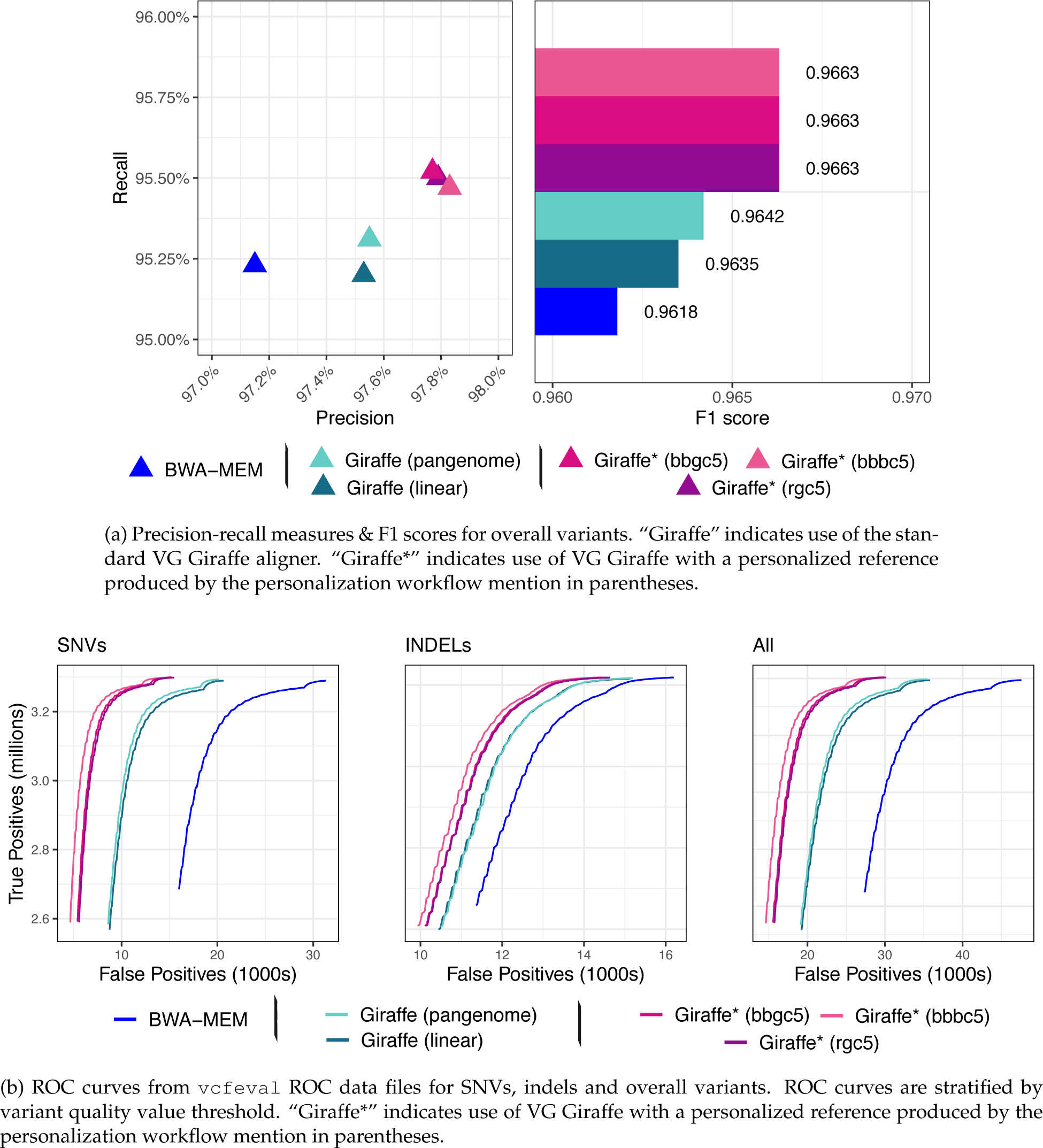
Variant calling accuracy metrics evaluated against T2TQ100 HG002 v1.0 truth VCF on GIAB HG002 high-confidence intersect regions. HG002 donor reads were aligned with a standard linear aligner (BWA-MEM) or one of various VG Giraffe workflows. “Giraffe” indicates use of the standard VG Giraffe aligner. “Giraffe*” indicates use of VG Giraffe with a personalized reference produced by the personalization workflow mention in parentheses.

## Supplementary Tables

**Table S1:** Alignment score comparison between the rgc1-imputed diploid personalized reference and the standard linear GRCh38 reference.

| **Alignment Score**<br>**Difference**<br>**(Personalized - GRCh38)** | **Read**<br>**Count** | **Fraction (%)** | **Mean** | **Median** | **Variance** | **Std. Deviation** |
| --- | --- | --- | --- | --- | --- | --- |
| Positive | 50,065,960 | 96.71% | 7.70 | 5.00 | 100.97 | 10.05 |
| Negative | 1,702,409 | 3.29% | -7.58 | -5.00 | 112.81 | 10.62 |

**Table S2:** Variant calling performance metrics for HG001 real donor reads, stratified by SNVs, indels, and overall variants, across different reference combinations within GIAB HG001 high-confidence regions.

| Mode | Workflow | Precision (%) | Recall (%) | F1 score |
| --- | --- | --- | --- | --- |
| All | BWA-MEM | 98.77 | 99.28 | 0.9902 |
|  | VG Giraffe (linear) | 99.17 | 99.24 | 0.9921 |
|  | VG Giraffe (pangenome) | 99.18 | 99.37 | 0.9928 |
|  | VG Giraffe* (rgc1) | 99.33 | 99.54 | 0.9943 |
|  | VG Giraffe* (rgc5) | 99.34 | 99.54 | 0.9944 |
|  | VG Giraffe* (bbgc5) | 99.34 | 99.56 | 0.9945 |
|  | VG Giraffe* (bbbc5) | 99.36 | 99.54 | 0.9945 |
| SNV | BWA-MEM | 98.77 | 99.29 | 0.9903 |
|  | VG Giraffe (linear) | 99.21 | 99.28 | 0.9925 |
|  | VG Giraffe (pangenome) | 99.22 | 99.41 | 0.9932 |
|  | VG Giraffe* (rgc1) | 99.37 | 99.60 | 0.9949 |
|  | VG Giraffe* (rgc5) | 99.38 | 99.60 | 0.9949 |
|  | VG Giraffe* (bbgc5) | 99.38 | 99.63 | 0.9951 |
|  | VG Giraffe* (bbbc5) | 99.41 | 99.60 | 0.9950 |
| Indel | BWA-MEM | 98.92 | 98.95 | 0.9894 |
|  | VG Giraffe (linear) | 99.06 | 98.80 | 0.9893 |
|  | VG Giraffe (pangenome) | 99.06 | 98.90 | 0.9898 |
|  | VG Giraffe* (rgc1) | 99.18 | 98.96 | 0.9907 |
|  | VG Giraffe* (rgc5) | 99.16 | 98.99 | 0.9908 |
|  | VG Giraffe* (bbgc5) | 99.18 | 98.97 | 0.9908 |
|  | VG Giraffe* (bbbc5) | 99.18 | 99.02 | 0.9910 |

**Table S3:** Variant calling performance metrics for HG001 real donor reads for overall variants, across different reference combinations within GIAB GRCh38 Complex Medically Relevant Gene (CMRG) regions.

| Workflow | Precision (%) | Recall (%) | F1 score |
| --- | --- | --- | --- |
| BWA-MEM | 85.37 | 97.87 | 0.9120 |
| VG Giraffe (linear) | 85.57 | 97.81 | 0.9128 |
| VG Giraffe (pangenome) | 85.58 | 98.03 | 0.9138 |
| VG Giraffe* (rgc1) | 85.55 | 98.22 | 0.9144 |
| VG Giraffe* (rgc5) | 85.57 | 98.21 | 0.9146 |
| VG Giraffe* (bbgc5) | 85.60 | 98.24 | 0.9148 |
| VG Giraffe* (bbbc5) | 85.77 | 98.19 | 0.9156 |

**Table S4:** Variant calling performance metrics for HG001 real donor reads for overall variants, across different reference combinations within GIAB GRCh38 stratifications.

| Region | Workflow | Precision (%) | Recall (%) | F1 score |
| --- | --- | --- | --- | --- |
| MHC | BWA-MEM | 81.13 | 92.52 | 0.8646 |
|  | VG Giraffe (linear) | 83.13 | 92.28 | 0.8746 |
|  | VG Giraffe (pangenome) | 82.03 | 93.02 | 0.8718 |
|  | VG Giraffe* (rgc1) | 84.63 | 92.59 | 0.8843 |
|  | VG Giraffe* (rgc5) | 84.90 | 92.58 | 0.8858 |
|  | VG Giraffe* (bbgc5) | 84.72 | 92.99 | 0.8867 |
|  | VG Giraffe* (bbbc5) | 88.77 | 91.80 | 0.9026 |
| alldifficultregions | BWA-MEM | 56.08 | 91.45 | 0.6952 |
|  | VG Giraffe (linear) | 59.21 | 91.72 | 0.7196 |
|  | VG Giraffe (pangenome) | 59.39 | 92.13 | 0.7222 |
|  | VG Giraffe* (rgc1) | 59.99 | 93.15 | 0.7298 |
|  | VG Giraffe* (rgc5) | 60.04 | 93.13 | 0.7301 |
|  | VG Giraffe* (bbgc5) | 60.00 | 93.28 | 0.7303 |
|  | VG Giraffe* (bbbc5) | 60.25 | 93.17 | 0.7318 |
| allOtherDifficultregions | BWA-MEM | 18.08 | 82.75 | 0.2967 |
|  | VG Giraffe (linear) | 23.28 | 82.59 | 0.3633 |
|  | VG Giraffe (pangenome) | 23.20 | 84.37 | 0.3639 |
|  | VG Giraffe* (rgc1) | 24.65 | 85.32 | 0.3824 |
|  | VG Giraffe* (rgc5) | 24.71 | 85.28 | 0.3832 |
|  | VG Giraffe* (bbgc5) | 24.70 | 85.62 | 0.3834 |
|  | VG Giraffe* (bbbc5) | 24.98 | 84.25 | 0.3854 |
| alllowmapandsegdupregions | BWA-MEM | 40.27 | 81.74 | 0.5395 |
|  | VG Giraffe (linear) | 47.10 | 83.40 | 0.6020 |
|  | VG Giraffe (pangenome) | 48.27 | 84.21 | 0.6137 |
|  | VG Giraffe* (rgc1) | 49.98 | 88.00 | 0.6375 |
|  | VG Giraffe* (rgc5) | 50.06 | 87.90 | 0.6379 |
|  | VG Giraffe* (bbgc5) | 50.03 | 88.46 | 0.6391 |
|  | VG Giraffe* (bbbc5) | 50.33 | 88.26 | 0.6410 |

**Table S5:** Time and memory taken by each VG Giraffe-based workflow in the personalization, downstream indexing, and downstream alignment steps.

| Phase | Workflow | Time (min) | Memory (GB) |
| --- | --- | --- | --- |
| Personalization | Impute-first: rgc1 | 50.23 | 14.45 |
|  | Impute-first: rgc5 | 94.97 | 14.45 |
|  | Impute-first: bbgc5 | 298.36 | 3.44 |
|  | Impute-first: bbbc5 | 329.17 | 34.78 |
| Giraffe autoindex | linear | 76.735 | 84.366 |
|  | pangenome | 1817.522 | 397.303 |
|  | Impute-first: rgc1 | 86.483 | 113.902 |
|  | Impute-first: rgc5 | 91.604 | 114.077 |
|  | Impute-first: bbgc5 | 91.910 | 114.783 |
|  | Impute-first: bbbc5 | 87.335 | 115.220 |
| Giraffe alignment | linear | 137.529 | 53.062 |
|  | pangenome | 204.824 | 70.086 |
|  | Impute-first: rgc1 | 157.973 | 65.249 |
|  | Impute-first: rgc5 | 158.795 | 63.799 |
|  | Impute-first: bbgc5 | 167.117 | 65.432 |
|  | Impute-first: bbbc5 | 160.250 | 65.440 |

**Table S6:** Index and graph size measurements for the VG Giraffe graphs generated in various workflows.

| Input to autoindex | .gbz<br>file (GB) | .dist<br>file (GB) | min<br>file (GB) | Total<br>(GB) | Size<br>(nodes + edges) |
| --- | --- | --- | --- | --- | --- |
| Linear GRCh38 | 2.6 | 1.5 | 35 | 39.1 | 193,945,149 |
| 1000 Genomes phase-3 pangenome | 5.4 | 4.8 | 35 | 45.2 | 469,910,808 |
| rgc1-imputed diploid | 3.1 | 2.3 | 35 | 40.4 | 255,676,786 |
| rgc5-imputed diploid | 3.1 | 2.3 | 35 | 40.4 | 255,645,758 |
| bbgc5-imputed diploid | 3.1 | 2.3 | 35 | 40.4 | 256,688,029 |
| bbbc5-imputed diploid | 3.1 | 2.3 | 35 | 40.4 | 258,286,403 |

**Table S7:** Variant calling performance metrics for all variants present in HG001 across different minor allele frequency (MAF) ranges.

| Variant Frequency | Workflow | Precision (%) | Recall (%) | F1 score |
| --- | --- | --- | --- | --- |
| MAF ≥ 5%<br>(n = 1,164,564) | BWA-MEM | 99.77 | 99.67 | 0.9972 |
|  | VG Giraffe (linear) | 99.81 | 99.69 | 0.9975 |
|  | VG Giraffe (pangenome) | 99.84 | 99.78 | 0.9981 |
|  | VG Giraffe* (rgc1) | 99.86 | 99.78 | 0.9982 |
|  | VG Giraffe* (rgc5) | 99.86 | 99.78 | 0.9982 |
|  | VG Giraffe* (bbgc5) | 99.86 | 99.79 | 0.9982 |
|  | VG Giraffe* (bbbc5) | 99.86 | 99.78 | 0.9982 |
| 0.5% ≤ MAF < 5%<br>(n = 171,987) | BWA-MEM | 99.96 | 99.87 | 0.9992 |
|  | VG Giraffe (linear) | 99.96 | 99.89 | 0.9993 |
|  | VG Giraffe (pangenome) | 99.97 | 99.92 | 0.9994 |
|  | VG Giraffe* (rgc1) | 99.97 | 99.92 | 0.9995 |
|  | VG Giraffe* (rgc5) | 99.97 | 99.93 | 0.9995 |
|  | VG Giraffe* (bbgc5) | 99.97 | 99.93 | 0.9995 |
|  | VG Giraffe* (bbbc5) | 99.97 | 99.92 | 0.9995 |
| MAF < 0.5%<br>(n = 58,503) | BWA-MEM | 99.98 | 99.71 | 0.9984 |
|  | VG Giraffe (linear) | 99.99 | 99.76 | 0.9988 |
|  | VG Giraffe (pangenome) | 99.99 | 99.78 | 0.9988 |
|  | VG Giraffe* (rgc1) | 99.99 | 99.84 | 0.9992 |
|  | VG Giraffe* (rgc5) | 99.99 | 99.85 | 0.9992 |
|  | VG Giraffe* (bbgc5) | 99.99 | 99.83 | 0.9991 |
|  | VG Giraffe* (bbbc5) | 99.99 | 99.83 | 0.9991 |

**Table S8:** Variant calling performance metrics for HG002 real donor reads, stratified by SNVs, indels, and overall variants, across different reference combinations using GIAB HG002 v4.2.1 truth within high-confidence regions.

| Mode | Workflow | Precision (%) | Recall (%) | F1 score |
| --- | --- | --- | --- | --- |
| All | BWA-MEM | 98.77 | 99.17 | 0.9897 |
|  | VG Giraffe (linear) | 99.24 | 99.09 | 0.9917 |
|  | VG Giraffe (pangenome) | 99.24 | 99.22 | 0.9923 |
|  | VG Giraffe* (rgc5) | 99.41 | 99.38 | 0.9939 |
|  | VG Giraffe* (bbgc5) | 99.42 | 99.41 | 0.9941 |
|  | VG Giraffe* (bbbc5) | 99.46 | 99.35 | 0.9941 |
| SNV | BWA-MEM | 98.77 | 99.20 | 0.9899 |
|  | VG Giraffe (linear) | 99.28 | 99.14 | 0.9921 |
|  | VG Giraffe (pangenome) | 99.27 | 99.27 | 0.9927 |
|  | VG Giraffe* (rgc5) | 99.45 | 99.44 | 0.9945 |
|  | VG Giraffe* (bbgc5) | 99.46 | 99.48 | 0.9947 |
|  | VG Giraffe* (bbbc5) | 99.51 | 99.41 | 0.9946 |
| Indel | BWA-MEM | 98.89 | 98.85 | 0.9887 |
|  | VG Giraffe (linear) | 99.11 | 98.71 | 0.9891 |
|  | VG Giraffe (pangenome) | 99.12 | 98.76 | 0.9894 |
|  | VG Giraffe* (rgc5) | 99.22 | 98.86 | 0.9904 |
|  | VG Giraffe* (bbgc5) | 99.26 | 98.83 | 0.9905 |
|  | VG Giraffe* (bbbc5) | 99.29 | 98.86 | 0.9908 |

**Table S9:** Variant calling performance metrics for HG002 real donor reads, stratified by SNVs, indels, and overall variants, across different reference combinations using T2TQ100 HG002 v1.0 truth within GIAB HG002 high-confidence regions.

| Mode | Workflow | Precision (%) | Recall (%) | F1 score |
| --- | --- | --- | --- | --- |
| All | BWA-MEM | 98.77 | 98.68 | 0.9872 |
|  | VG Giraffe (linear) | 99.07 | 98.67 | 0.9887 |
|  | VG Giraffe (pangenome) | 99.08 | 98.77 | 0.9892 |
|  | VG Giraffe* (rgc5) | 99.22 | 98.92 | 0.9907 |
|  | VG Giraffe* (bbgc5) | 99.23 | 98.94 | 0.9908 |
|  | VG Giraffe* (bbbc5) | 99.25 | 98.90 | 0.9907 |
| SNV | BWA-MEM | 99.07 | 99.29 | 0.9918 |
|  | VG Giraffe (linear) | 99.39 | 99.29 | 0.9934 |
|  | VG Giraffe (pangenome) | 99.40 | 99.40 | 0.9940 |
|  | VG Giraffe* (rgc5) | 99.55 | 99.57 | 0.9956 |
|  | VG Giraffe* (bbgc5) | 99.56 | 99.57 | 0.9957 |
|  | VG Giraffe* (bbbc5) | 99.58 | 99.54 | 0.9956 |
| Indel | BWA-MEM | 96.92 | 94.93 | 0.9592 |
|  | VG Giraffe (linear) | 97.11 | 94.79 | 0.9594 |
|  | VG Giraffe (pangenome) | 97.12 | 94.84 | 0.9597 |
|  | VG Giraffe* (rgc5) | 97.24 | 94.86 | 0.9604 |
|  | VG Giraffe* (bbgc5) | 97.25 | 94.87 | 0.9605 |
|  | VG Giraffe* (bbbc5) | 97.25 | 94.93 | 0.9608 |

**Table S10:** Variant calling performance metrics for HG002 real donor reads, stratified by SNVs, indels, and overall variants, across different reference combinations using T2TQ100 HG002 v1.0 truth within T2TQ100 HG002 v1.0 high-confidence regions.

| Mode | Workflow | Precision (%) | Sensitivity (%) | F1 score |
| --- | --- | --- | --- | --- |
| All | BWA-MEM | 96.94 | 95.09 | 96.01 |
|  | VG Giraffe (linear) | 97.39 | 95.03 | 96.19 |
|  | VG Giraffe (pangenome) | 97.40 | 95.15 | 96.26 |
|  | VG Giraffe* (rgc5) | 97.63 | 95.35 | 96.48 |
|  | VG Giraffe* (bbgc5) | 97.61 | 95.38 | 96.48 |
|  | VG Giraffe* (bbbc5) | 97.67 | 95.33 | 96.49 |
| SNP | BWA-MEM | 98.15 | 97.88 | 98.02 |
|  | VG Giraffe (linear) | 98.64 | 97.90 | 98.27 |
|  | VG Giraffe (pangenome) | 98.66 | 98.02 | 98.34 |
|  | VG Giraffe* (rgc5) | 98.90 | 98.27 | 98.59 |
|  | VG Giraffe* (bbgc5) | 98.90 | 98.27 | 98.59 |
|  | VG Giraffe* (bbbc5) | 98.97 | 98.20 | 98.59 |
| Indel | BWA-MEM | 92.77 | 84.31 | 88.34 |
|  | VG Giraffe (linear) | 92.99 | 83.99 | 88.26 |
|  | VG Giraffe (pangenome) | 92.97 | 84.07 | 88.30 |
|  | VG Giraffe* (rgc5) | 93.12 | 84.16 | 88.42 |
|  | VG Giraffe* (bbgc5) | 93.27 | 84.04 | 88.42 |
|  | VG Giraffe* (bbbc5) | 93.17 | 84.18 | 88.45 |

**Table S11:** Variant calling performance metrics for HG002 real donor reads for overall variants, across different reference combinations within GIAB GRCh38 Complex Medically Relevant Gene (CMRG) regions.

| **Workflow** | **Precision (%)** | **Recall (%)** | **F1 score** |
| --- | --- | --- | --- |
| BWA-MEM | 99.16 | 99.38 | 0.9927 |
| VG Giraffe (linear) | 99.39 | 99.33 | 0.9936 |
| VG Giraffe (pangenome) | 99.41 | 99.45 | 0.9943 |
| VG Giraffe* (rgc5) | 99.51 | 99.58 | 0.9954 |
| VG Giraffe* (bbgc5) | 99.48 | 99.59 | 0.9954 |
| VG Giraffe* (bbbc5) | 99.60 | 99.54 | 0.9957 |

**Table S12:** Variant calling performance metrics for HG002 real donor reads for overall variants, across different reference combinations within GIAB GRCh38 stratifications.

| Region | Workflow | Precision (%) | Recall (%) | F1 score |
| --- | --- | --- | --- | --- |
| MHC | BWA-MEM | 97.81 | 97.46 | 0.9764 |
|  | VG_Giraffe(linear) | 98.07 | 96.29 | 0.9717 |
|  | VG_Giraffe(pangenome) | 99.25 | 98.05 | 0.9865 |
|  | VG_Giraffe*(rgc5) | 99.46 | 98.19 | 0.9882 |
|  | VG_Giraffe*(bbgc5) | 99.45 | 97.95 | 0.9869 |
|  | VG_Giraffe*(bbbc5) | 98.67 | 94.06 | 0.9631 |
| alldifficultregions | BWA-MEM | 96.20 | 97.00 | 0.9660 |
|  | VG_Giraffe(linear) | 97.85 | 96.69 | 0.9727 |
|  | VG_Giraffe(pangenome) | 97.85 | 97.15 | 0.9750 |
|  | VG_Giraffe*(rgc5) | 98.32 | 97.75 | 0.9803 |
|  | VG_Giraffe*(bbgc5) | 98.35 | 97.87 | 0.9811 |
|  | VG_Giraffe*(bbbc5) | 98.51 | 97.67 | 0.9809 |
| allOtherDifficultregions | BWA-MEM | 72.17 | 90.37 | 0.8025 |
|  | VG_Giraffe(linear) | 83.83 | 89.58 | 0.8661 |
|  | VG_Giraffe(pangenome) | 84.23 | 90.80 | 0.8739 |
|  | VG_Giraffe*(rgc5) | 88.57 | 92.67 | 0.9057 |
|  | VG_Giraffe*(bbgc5) | 88.82 | 92.95 | 0.9084 |
|  | VG_Giraffe*(bbbc5) | 90.23 | 91.44 | 0.9083 |
| alllowmapandsegdupregions | BWA-MEM | 89.19 | 90.68 | 0.8993 |
|  | VG_Giraffe(linear) | 94.51 | 90.01 | 0.9221 |
|  | VG_Giraffe(pangenome) | 94.48 | 91.51 | 0.9297 |
|  | VG_Giraffe*(rgc5) | 95.93 | 93.71 | 0.9481 |
|  | VG_Giraffe*(bbgc5) | 96.08 | 94.18 | 0.9512 |
|  | VG_Giraffe*(bbbc5) | 96.52 | 93.55 | 0.9501 |

**Table S13:** Variant calling performance metrics for HG002 real donor reads for overall variants, across different reference combinations within GIAB HG002 v1.0 Complex Medically Relevant Gene (CMRG) high-confidence regions.

| Workflow | Precision (%) | Recall (%) | F1 score |
| --- | --- | --- | --- |
| BWA-MEM | 94.70 | 95.10 | 0.9490 |
| VG Giraffe (linear) | 95.84 | 94.39 | 0.9511 |
| VG Giraffe (pangenome) | 95.83 | 95.05 | 0.9544 |
| VG Giraffe* (rgc5) | 93.57 | 95.35 | 0.9445 |
| VG Giraffe* (bbgc5) | 93.86 | 95.33 | 0.9459 |
| VG Giraffe* (bbbc5) | 95.25 | 95.23 | 0.9524 |

**Table S14:** Variant calling performance metrics for all variants present in HG002 across different minor allele frequency (MAF) ranges.

| Variant Frequency | Workflow | Precision (%) | Recall (%) | F1 score |
| --- | --- | --- | --- | --- |
| MAF ≥ 5% (n=1,179,717) | BWA-MEM | 99.83 | 99.72 | 0.9977 |
|  | VG Giraffe (linear) | 99.86 | 99.76 | 0.9981 |
|  | VG Giraffe (pangenome) | 99.89 | 99.85 | 0.9987 |
|  | VG Giraffe* (rgc5) | 99.90 | 99.84 | 0.9987 |
|  | VG Giraffe* (bbgc5) | 99.90 | 99.85 | 0.9988 |
|  | VG Giraffe* (bbbc5) | 99.90 | 99.80 | 0.9985 |
| 0.5% ≤ MAF < 5% (n=175,103) | BWA-MEM | 99.97 | 99.88 | 0.9992 |
|  | VG Giraffe (linear) | 99.97 | 99.90 | 0.9994 |
|  | VG Giraffe (pangenome) | 99.98 | 99.93 | 0.9995 |
|  | VG Giraffe* (rgc5) | 99.98 | 99.93 | 0.9995 |
|  | VG Giraffe* (bbgc5) | 99.98 | 99.94 | 0.9996 |
|  | VG Giraffe* (bbbc5) | 99.99 | 99.90 | 0.9994 |
| MAF < 0.5% (n=60,273) | BWA-MEM | 99.97 | 99.73 | 0.9985 |
|  | VG Giraffe (linear) | 99.98 | 99.74 | 0.9986 |
|  | VG Giraffe (pangenome) | 99.98 | 99.77 | 0.9988 |
|  | VG Giraffe* (rgc5) | 99.99 | 99.83 | 0.9991 |
|  | VG Giraffe* (bbgc5) | 99.98 | 99.85 | 0.9992 |
|  | VG Giraffe* (bbbc5) | 99.98 | 99.84 | 0.9991 |

**Table S15:** Variant calling performance metrics for all variants present in HG002 CMRG v1.0 across different minor allele frequency (MAF) ranges.

| Variant Frequency | Workflow | Precision (%) | Recall (%) | F1 score |
| --- | --- | --- | --- | --- |
| MAF ≥ 5% (n=5,718) | BWA-MEM | 99.54 | 98.51 | 0.9902 |
|  | VG Giraffe (linear) | 99.60 | 98.24 | 0.9892 |
|  | VG Giraffe (pangenome) | 99.58 | 98.53 | 0.9905 |
|  | VG Giraffe* (rgc5) | 99.68 | 98.69 | 0.9918 |
|  | VG Giraffe* (bbgc5) | 99.66 | 98.81 | 0.9923 |
|  | VG Giraffe* (bbbc5) | 99.58 | 98.53 | 0.9905 |
| 0.5% ≤ MAF < 5% (n=692) | BWA-MEM | 99.68 | 99.21 | 0.9944 |
|  | VG Giraffe (linear) | 99.84 | 99.05 | 0.9944 |
|  | VG Giraffe (pangenome) | 99.84 | 99.21 | 0.9952 |
|  | VG Giraffe* (rgc5) | 100.00 | 99.21 | 0.9960 |
|  | VG Giraffe* (bbgc5) | 100.00 | 99.21 | 0.9960 |
|  | VG Giraffe* (bbbc5) | 99.84 | 99.21 | 0.9952 |
| MAF < 0.5% (n=245) | BWA-MEM | 98.88 | 92.67 | 0.9567 |
|  | VG Giraffe (linear) | 98.88 | 93.19 | 0.9595 |
|  | VG Giraffe (pangenome) | 98.88 | 93.19 | 0.9595 |
|  | VG Giraffe* (rgc5) | 100.00 | 95.81 | 0.9786 |
|  | VG Giraffe* (bbgc5) | 100.00 | 95.81 | 0.9786 |
|  | VG Giraffe* (bbbc5) | 98.89 | 93.72 | 0.9623 |

**Table S16:** Summary of rare HG002 CMRG v1.0 variants recovered by *Impute-first (rgc5, bbgc5)* workflows in the TPO gene region [ OMIM, gnomAD ].

| CHROM | POS | REF > ALT | dbSNP ID | gnomAD v4.1.0 ID<sup>†</sup> | Frequency* |
| --- | --- | --- | --- | --- | --- |
| chr2 | 1424939 | C > T | rs113311329 | 2-1424939-C-T | 0.001 |
| chr2 | 1424955 | T > C | rs541492938 | 2-1424955-T-C | 0.0006 |
| chr2 | 1424964 | A > G | rs563945547 | 2-1424964-A-G | 0.0014 |
| chr2 | 1424965 | A > C | rs531247861 | 2-1424965-A-C | 0.0012 |

\* dbSNP reported Variation Frequency (1000G).
<sup>†</sup> Rows 2& 3 are not observed in gnomAD records. All links show Variant Effect Predictor (VEP) annotation analysis. According to VEP annotation analysis, these variants fall on 10 transcripts in 2 genes suggesting a potential impact on gene function and possible clinical relevance.

